# Disease-associated microRNA, miR-9-2, regulates timing of retinal progenitor cell competence and maintenance of Müller glial identity

**DOI:** 10.64898/2026.03.27.714903

**Authors:** LuLu K Callies, Aarushi Jain, Stella Xu, Emma D Thomas, Timothy J Cherry

**Author notes:** CORRESPONDENCE (T.J.C.).

## Abstract

**ABSTRACT/SUMMARY:** Development of the visual system is dependent upon precise regulation of cell fate specification. In the mammalian retina, a single pool of multipotent progenitor cells becomes competent to produce the seven major retinal cell classes in distinct but overlapping windows. MicroRNAs (miRNAs) have been implicated in controlling retinal progenitor competence and risk for retinal disease, but the specific contribution of individual miRNAs and how they may be regulated is still unclear. Here we characterize a deeply conserved gene regulatory unit that includes the miRNA, miR-9-2, and a retinal-disease-associated enhancer that controls its expression. Loss of miR-9-2, one of three mammalian miR-9 paralogs, delays the emergence of late-born retinal cell classes and leads to misspecification of Müller glial cells to a hybrid neuronal-glial fate. Further, we identify transcription factors and gene regulatory networks directly controlled by miR-9-2 during retinal development. Lastly, we provide evidence of a negative feedback loop through which miR-9-2 regulates itself. Altogether, this study provides insight into mechanisms that regulate the timing of retinal progenitor competence and glial cell identity, and how this gene regulatory unit may contribute to retinal disease.

**HIGHLIGHTS:**

- A functionally conserved, disease-associated enhancer regulates miR9-2 expression in human and mouse retina.
- miR9-2 regulates key transcription factors in progenitor cells and glia.
- miR9-2 controls the timing of retinal cell class specification.
- Regulation of miR9-2 is required to establish and maintain proper glial cell identity.

## INTRODUCTION

Precise regulation of cell fate specification is essential for generating a functional central nervous system (CNS). In the mammalian retina, a single pool of retinal progenitor cells (RPCs) gives rise to all seven major retinal cell classes with a precise order and timing. These RPCs become competent to produce different cell classes in distinct but overlapping windows. In the retina, cell class birth is often split into two phases: early-born neurons – retinal ganglion (RGCs), cone photoreceptor, and horizontal cells (HCs) – produced first, followed by the late-born neurons – rod photoreceptor and bipolar cells (BCs) – and the primary glial cell class of the retina, Müller glia (MG)^1–4^. The amacrine cell class (ACs) spans these two developmental epochs with early-born ACs, like cholinergic starburst amacrine cells, and later-born ACs, such as the glycinergic amacrine cell types.

The genetic programs that influence retinal cell class specification are well studied. Transcription factors (TFs) have documented roles in modulating progenitor cell competence. Among them, basic helix-loop-helix proteins (bHLHs), homeodomain-containing TFs, Notch signaling proteins, and Nuclear Factor I (NFI) family members have been found to play roles in regulating cell proliferation, cell cycle exit, and cell class specification in the retina. TFs, such as the bHLH and Onecut genes, have documented roles in promoting specification of early-born cell types, including RGC fate (Atoh7^3,5–7)^ or horizontal and cone cell fate (Onecuts^8–13)^. Conversely, Notch signaling proteins and NFI factors are thought to maintain progenitor identity. Notch pathway signaling early in development, through effectors such as *Hes1* and *Hes5*, maintain RPCs in a proliferative state^14–18^, while recent work on NFI factors found that they regulate specification of late-born cell fates, including bipolar and MG cells^19^. While the roles of these key TFs are well understood, how the factors themselves are regulated across development allowing for robust transitions in cell fate specification is not fully understood.

MicroRNAs (miRNAs) have been implicated in controlling the timing of progenitor cell competence in the CNS. miRNAs are a class of small, single-stranded RNAs around 20 nucleotides in length and act as translational repressors, binding in a complementary fashion to target seed sequences in the 3’ UTR of mRNAs which leads to repression and degradation of the transcript^20,21^. Many miRNAs arose from ancient gene duplication events and are classified into families composed of miRNA paralogs which target the same seed sequences in their processed form but originate from different host genes and evolved to have different patterns of expression in distinct cell types^21^. In the mammalian retina, the importance of miRNAs in regulating retinal progenitor competence and homeostasis is clear, as deletion of Dicer, an enzyme important to processing miRNAs into their mature form, leads to a delay in the production of late-born cell types^22–25^. Further, the miR-9 family, among others, was identified as a potential regulator of the shift in competence from early- to late-born cell types due to its increased expression when RPCs undergo this transition.

The highly-conserved miR-9 family of microRNAs has previously been implicated in neural development in diverse model organisms^20,26^. Studies investigating neuronal miR-9 function in *Drosophila,* zebrafish and mouse telencephalon, have found that loss of this miRNA can lead to increased progenitor proliferation and excess neuronal precursors, delayed cell cycle exit, and/or abnormal vascularization of CNS tissues, including the retina^20,27–30^. Mechanistically, these studies have proposed candidate miR-9 targets including *Hes1*, thereby influencing Notch signaling, as well as Onecut and Tlx TFs^20,28,29,31,32^. In the mammalian retina, overexpression of miR-9 seems to bias cells towards neuronal rather than glial fates^33^. While these studies have proposed many functions for this microRNA family, there is still a significant gap in knowledge about the roles of individual miR-9 family members, especially as each paralog displays distinct tissue and cell type-specific expression. In mice and humans, there are three miR-9 paralogs, known as miR-9-1, 2, and 3^20^. Though prior studies implicate miR-9 as a regulator of neurogenesis, the specific targets of the microRNA are not well defined, and approaches thus far have more broadly disrupted miRNA function rather than focusing on the behavior of specific miR-9 family members in specific cell types.

Genome wide association studies (GWAS) have increasingly linked the miR-9-2 host gene locus to retinal diseases, such as Macular Telangiectasia Type II and Age-Related Macular Degeneration, and disease phenotypes, including altered retinal vasculature caliber and altered retinal thickness^34–39^. More specifically, an enhancer for miR-9-2, recently investigated by our lab, has been implicated in these GWAS (Fig. 1A). Using single-nucleus Assay for Transposase-Accessible Chromatin sequencing (snATAC-seq), we found that the miR-9-2 enhancer is accessible in RPCs and MG of the human retina. Further, we determined that, in human retinal organoids, knockout of this enhancer leads to a decrease in miR-9-2 host gene expression and aberrant developmental phenotypes, including altered cell class proportions indicative of a delay in the emergence of various cell classes^40^. These data suggests that miR-9-2, independent of miR-9-1 and miR-9-3, can play an important regulatory role in the retina. This work improved our understanding of the functional significance of miR-9-2 *in vitro* in developing human tissue, but there are still many questions remaining surrounding how miR-9-2 functions *in vivo* to regulate retinal development and disease. Our work on the miR-9-2 enhancer, alongside pre-existing studies on the miR-9 family, indicates that miR-9-2 may be important in RPCs for properly specifying cell types and controlling the relative abundance of types in the mature retina. Additionally, the strong linkage of miR-9-2 with retinal disease and macular degenerative disorders suggests that miR-9-2 is important to maintaining glial homeostasis and preventing the MG cell loss that is frequently found in retinal degenerative diseases. These outstanding questions required an *in vivo* system to study miR-9-2 in the developing and mature retina.

**Figure 1.**
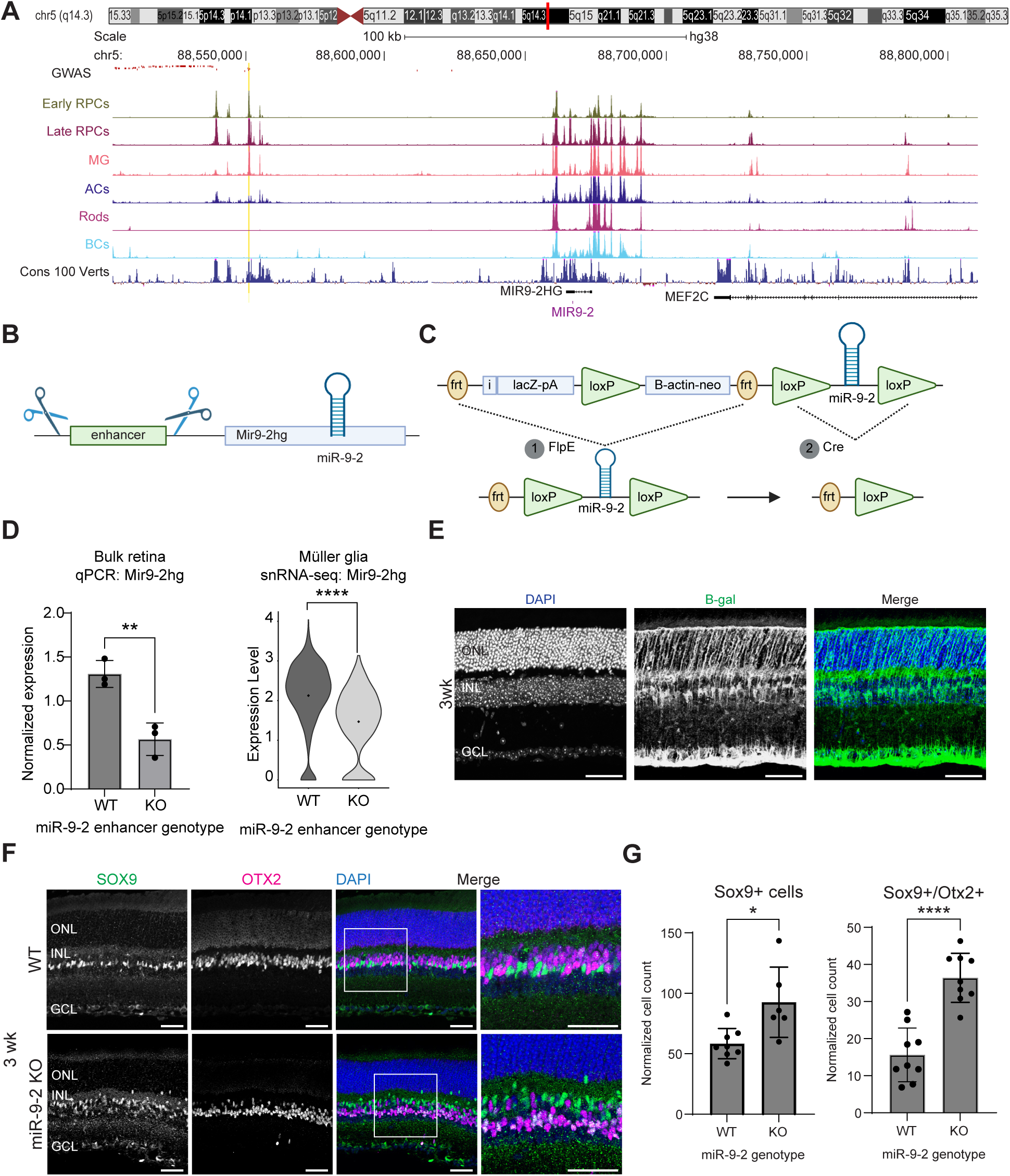
Disruption to disease-associated miR-9-2 regulatory locus leads to glial misspecification in developing mouse retina. **A.** UCSC human genome browser tracks (hg38) of the genomic location of the microRNA, miR-9-2 and its host gene. GWAS variants associated with Macular Telangiectasia Type 2 (MacTel) are depicted as red lines^34^ with the lead risk variant highlighted in yellow. Tracks show accessibility from single-nucleus ATAC-sequencing of human retina (all scaled 0-55) with bottom track showing regions conserved across vertebrate species^40^. **B.** Schematic of enhancer deletion by CRISPR targeting to generate miR-9-2 enhancer knockout mouse. **C.** Schematic of generation of germline miR-9-2 KO mice. **D.** qPCR of bulk retinal tissue (n=3) and snRNA-seq of MG (n=3) comparing mean *Mir9-2-hg* expression between miR-9-2 enhancer knockout and WT retinas. qPCR: mean +/- 1SD, student’s t-test, **p≤0.01. snRNA-seq: mean represented as point, Wilcoxon rank sum test with Benjamini-Hochberg correction, ****p≤0.0001. **E.** Immunolabeling of β-galactosidase (green) to visualize expression of lacZ as proxy for miR-9-2 expression in 3-week mice containing floxed miR-9-2 hairpin and β-gal reporter cassette. **F.** Immunolabeling for MG (SOX9, green) and BCs (OTX2, magenta) in cross-sections of WT and KO retinas. Cells co-expressing OTX2 and SOX9 indicated by white arrowheads. **G.** Cell count values normalized to retinal area for SOX9+ (WT, n=7; KO, n=6) and SOX9+/OTX2+ (WT, n=9; KO, n=9) cells in KO and WT retinas. Counts are average of two images of the same retina in corresponding regions. Values reported as mean +/- 1SD. Mann Whitney test, *p≤0.05, ****p≤0.0001. All scale bars = 50 microns. Abbreviations: RPC = retinal progenitor cell, MG = Müller glia, AC = amacrine cell, BC = bipolar cell, ONL = outer nuclear layer, INL = inner nuclear layer, GCL = ganglion cell layer

In this study, we sought to investigate the regulatory role of miR-9-2 in the specification and maintenance of retinal cell identity. Using novel enhancer knockout and miR-9-2 germline knockout (KO) mouse models, immunohistochemistry, and single-nucleus RNA-sequencing (snRNA-seq), we found miR-9-2 regulates the timing of retinal cell fate emergence. Developing miR-9-2 KO mouse retinas contain larger proportions of early-born cell classes compared to sex-matched, littermate controls. We also found that miR-9-2 is necessary at the end of development for MG to adopt and maintain their mature glial cell fate. In enhancer and miR9-2 knockout retinas, MG are mis-specified adopting hybrid neuronal- and glial-like cell fates. Further, we identify direct targets of miR-9-2 in developing and mature cell populations and the downstream gene regulatory networks (GRNs) that influence cell fate. This work expands our knowledge of how the temporal dynamics of retinal cell class specification are controlled during development and adds to our understanding of miR-9-2 in retinal disease risk.

## RESULTS

### Conserved regulation of a retinal disease-associated microRNA

To examine miR-9-2 regulation and function *in vivo*, we generated a germline, enhancer knockout mouse model by CRISPR targeting a conserved ∼300bp region corresponding to accessibility (snATAC-seq data) in the putative enhancer region (Fig. 1A-B). Retinas from adult animals were used for qPCR analysis on bulk retinal tissue (n=3) targeting the miR-9-2 host gene (*Mir9-2hg*). Consistent with our human organoid studies, we found a significant decrease of the *Mir9-2hg* expression in enhancer knockout animals compared to WT (Fig. 1D). Using single-nuclei RNA-sequencing (snRNA-seq; n=3), we also profiled WT and enhancer knockout retinas from mature animals to understand enhancer activity in a cell-specific manner (SFig. 1). We found that MG from enhancer knockout animals had a significant decrease in *Mir9-2hg* expression compared to WT (Fig. 1D; SFig. 1E). We also identified other significant differentially expressed genes (DEGs), particularly in MG cells, several of which are computationally predicted to be direct miR-9 targets like Foxp2 (SFig. 1E). These results demonstrate that this 5q14.3 enhancer has a conserved role in regulating the miR-9-2 host gene, and thereby miR-9-2, across humans and mice.

Our previous work demonstrated that knockout of the miR-9-2 enhancer in retinal organoids led to defects in retinal development^40^. To determine whether the miR-9-2 enhancer has a similar role in mice, we analyzed cell class percentages from the snRNA-seq data and did not observe any significant differences (SFig. 1D). To validate this and determine if there were more subtle morphological differences, we performed immunolabeling for major markers of retinal cell classes in mature enhancer knockout and WT retinas. We did not observe any significant changes in cell class numbers (SFig. 1D, 1F; SFig. 2). However, we did observe a change in the positioning of Sox9 positive (+) cells, where Sox9+ KO cells appeared to have migrated towards the choroidal side of the retina and were more interspersed with Otx2+ BCs (SFig. 2A).

To more directly dissect the role of miR-9-2 in retinal development, we developed a germline miR-9-2 hairpin KO mouse model. First, taking advantage of a previously published strain of *Mir-9-2^tm1Mtm^* mice (JAX:036062-JAX^41^) containing a knocked-in floxed allele of the miR-9-2 hairpin loop and an upstream lacZ reporter cassette surrounded by Frt sites, we assessed lacZ expression as a proxy for miR-9-2 transcript expression. By immunolabelling ß-galactosidase (ß-gal) in 5-week-old mouse retinas, we visualized which cell classes expressed the reporter as proxy for miR-9-2. In alignment with our previously reported accessibility and expression of the *MIR9-2HG* in human retina and human retinal organoids^40^, we confirmed MG-specific expression of miR-9-2 in the adult mouse retina. ß-gal+ cells had morphological features of MG cells, with long projections extending apically and basally through the retina and nuclei positioned within the inner nuclear layer (INL; Figure 1E). There did not appear to be other cell classes with reporter expression in the mature retina.

To generate the constitutive miR-9-2 knockout (KO), we crossed *Mir9-2^tm1Mtm^*mice to transgenic Rosa26::FLPE knock-in mice (JAX:009086^42^) to remove the lacZ reporter cassette and crossed the resulting progeny to E2a-cre mice (JAX:003724^43^), which express germline Cre, to remove miR-9-2 from all cells. Notably, only the miR-9-2 hairpin loop and nearby flanking sequence are deleted, while the remainder of the host gene is intact and able to be transcribed. We were able to confirm this directly by snRNA-seq, where we see that the host gene continues to be expressed even in KO retinas (SFig. 3).

To determine if cell fate specification was altered in miR-9-2 KO animals, we performed immunolabeling for common markers of the major cell classes of the retina in 3-week WT and KO animals. Notably, in KO retinas, we found an increase in SOX9+ cells, a marker for retinal Müller cells (Fig. 1F-G). Reminiscent of the enhancer knockout retinas, SOX9+ cells also no longer maintained normal positioning within the retina. In the wildtype INL, while the nuclei of SOX9+ MG typically reside below the nuclei of OTX2+ BCs (vitreal), in the KO retinas, many of the SOX9+ cells appeared to have migrated above the OTX2+ BCs (choroidal). Several of the KO MG cells also entered the outer nuclear layer (ONL) which typically contains exclusively photoreceptor nuclei (Fig. 1F). Surprisingly, many of the knockout SOX9+cells co-expressed OTX2, a BC marker (Fig. 1F-G). These data indicate altered gliogenesis in miR-9-2 KO retinas, leading to non-canonical marker expression and misspecification.

### Delayed cell class emergence in miR-9-2 KO retinas revealed by snRNA-Seq

To understand the origins of glial mispositioning and misspecification in the mature retina, we took advantage of snRNA-seq profiling and immunofluorescence. Normal retinal development occurs in a highly stereotyped manner, with a well-defined order and timing for the emergence of different cell classes (Fig. 2A). Prior literature suggests an important role for microRNAs during the transition from early- to late-born cell classes^22^. To determine the timing of miR-9 family member expression, we used a published single-cell atlas of mouse retinal development^19^ and found that expression of miR-9 host genes begins increasing at E16.5, peaking from E18.5-P5 (Fig. 2B). Further, *Mir9-2hg* appears to be the predominant miR-9 family member during the entire time-course of retinal development, emphasizing its unique importance.

**Figure 2.**
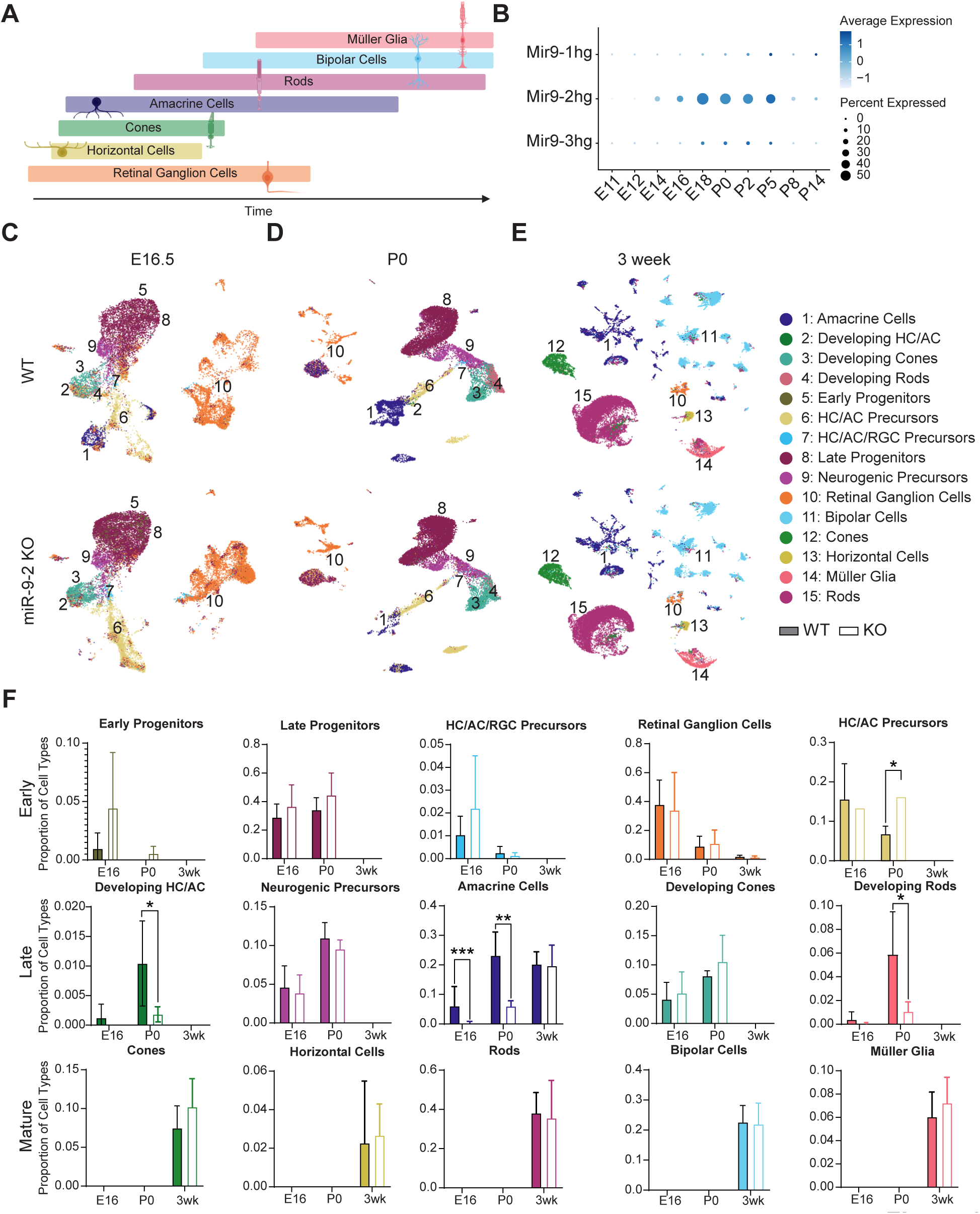
Loss of miR-9-2 disrupts retinal cell class proportions during retinal development. **A.** Schematic of overlapping windows of retinal cell class birth. **B.** Dot plot of average expression of miR-9 host gene paralogs in mouse retina over time, from published retinal single-cell RNA-sequencing atlas^19^. **C-E.** UMAPs of annotated cell classes in miR-9-2 WT and KO mice at E16.5 **(C)**, P0 **(D)**, and 3-weeks **(E)**. **F**. Cell class proportions comparing WT (solid bar) and miR-9-2 KO (empty bar) retinas for individual cell classes over time. Mean +/- 1SD. Significance of WT to KO comparison was determined using EdgeR generalized linear model with Benjamini-Hochberg correction^88,91^ *p≤0.05, **p≤0.01, ***p≤0.001. Abbreviations: RGC = retinal ganglion cells, HC = horizontal cells, AC = amacrine cells, BC = bipolar cells, MG = Müller glia.

Next, we performed snRNA-seq on miR-9-2 KO and sex-matched littermate control retinas at key timepoints during retinal development. We profiled WT and miR-9-2 KO retinas at E16.5 to capture the early-to-late transitional period, P0 to profile during the window of peak late-born cell birth, and adult retinas at 3-weeks to assess the end of development (E16.5 & P0, n=4; 3-weeks, n=5; per genotype). At all timepoints, we identified and annotated the developmentally expected cell classes (SFig. 4). We then assessed the relative proportions of cell classes between WT and KO retinas to determine whether cell classes emerge normally in the absence of miR-9-2. We found that direct loss of miR-9-2 *in vivo* perturbs normal cell class proportions (Table S1). In developing retinas, the proportions of differentiated ACs (E16.5, P0) and developing HC/ACs (P0 only) were significantly decreased in KO retinas compared to WT (Fig. 2C-D, 2F; SFig. 5A). Correspondingly, there was a significant increase in the proportions of HC/AC Precursors (P0 only) in the KO compared to WT (Figure 2D, 2F). This increase in the precursor populations and decrease in developing and mature cells indicates that loss of miR-9-2 leads to a more immature amacrine population.

Dramatically, the emergence of rod photoreceptors, the most abundant cell class, was also disrupted. At P0, there was a significantly smaller proportion of developing rods in miR9-2 KO retinas compared to the WT. Newborn miR-9-2 KO retinas appeared to almost completely lack rods (Fig. 2D, 2F). To determine whether this lack of photoreceptors could be attributed to cell death, we used Terminal deoxynucleotidyl transferase dUTP Nick-End Labeling (TUNEL) and immunolabeling for Phospho-Caspase 3 but found no differences between WT and KO animals (SFig. 6). As rods are the dominant late-born cell type (Fig. 2A), this suggests that rods are not yet born in miR-9-2 KO retinas, rather than dying. This phenotype is similar to prior studies which found that loss of Dicer in the retina leads to an inability of RPCs to produce late-born cell types^22–24^.

Despite these developmental differences, by 3-weeks, there were no significant differences in cell class proportions between WT and miR-9-2 KO retinas (Fig. 2E-F). While proportions no longer differed, we know MG still developed abnormal phenotypes, including migration towards the ONL and mis-expression of bipolar markers (Fig. 1E-F). This suggests that the major cell classes or unique cell types of the retina may still have altered transcriptional profiles, even as cell proportions are unchanged. Overall, these preliminary snRNA-seq analyses suggest that differences between WT and KO retinas during development are due to a delay in cell class production rather than a loss of late-born cells, like rods. Therefore, while it seems that miR-9-2 is important for the proper timing of cell class emergence, retinas lacking miR-9-2 still retain the ability to produce the major cell types, albeit with disrupted Müller glia.

### miR-9-2 is dynamically expressed in retinal progenitor cells and Müller glia

To understand more fully how loss of miR-9-2 alters the timing of cell class emergence and cell identity, we sought to identify which cell classes miR-9 is expressed in. Prior snATAC-seq data from our lab^40^ found that the enhancer of *MIR9-2HG* was most accessible in RPCs and MG in the developing and mature human retina, and at corresponding timepoints in human retinal organoids (Figure 1A, previously published work^40^). To determine whether this cell type-specific activity was recapitulated in the mouse retina, we used our snRNA-seq dataset to examine the expression levels of *Mir9-2hg* across cell types at each profiled timepoint. Using expression profiles from our WT mice, we found that at both E16.5 and P0, early and late RPCs had the highest *Mir9-2hg* expression, followed by several populations of precursor cells. In the mature, 3-week-old mouse retina, *Mir9-2hg* was predominantly expressed in MG. Further, as there are three paralogs of miR-9 in the mouse, we checked the expression of miR-9-1 and miR-9-3 host genes; at developmental timepoints, *Mir9-2hg* was the most predominantly expressed miR-9 host gene in RPCs (Figure 3A, 3C; SFig. 7). At 3-weeks, the expression of miR-9 paralogs was highly specific to MG cells. While *Mir9-2hg* was still the most predominant paralog expressed, *Mir9-1hg* expression was also detectable (Fig. 3E; SFig. 7).

**Figure 3.**
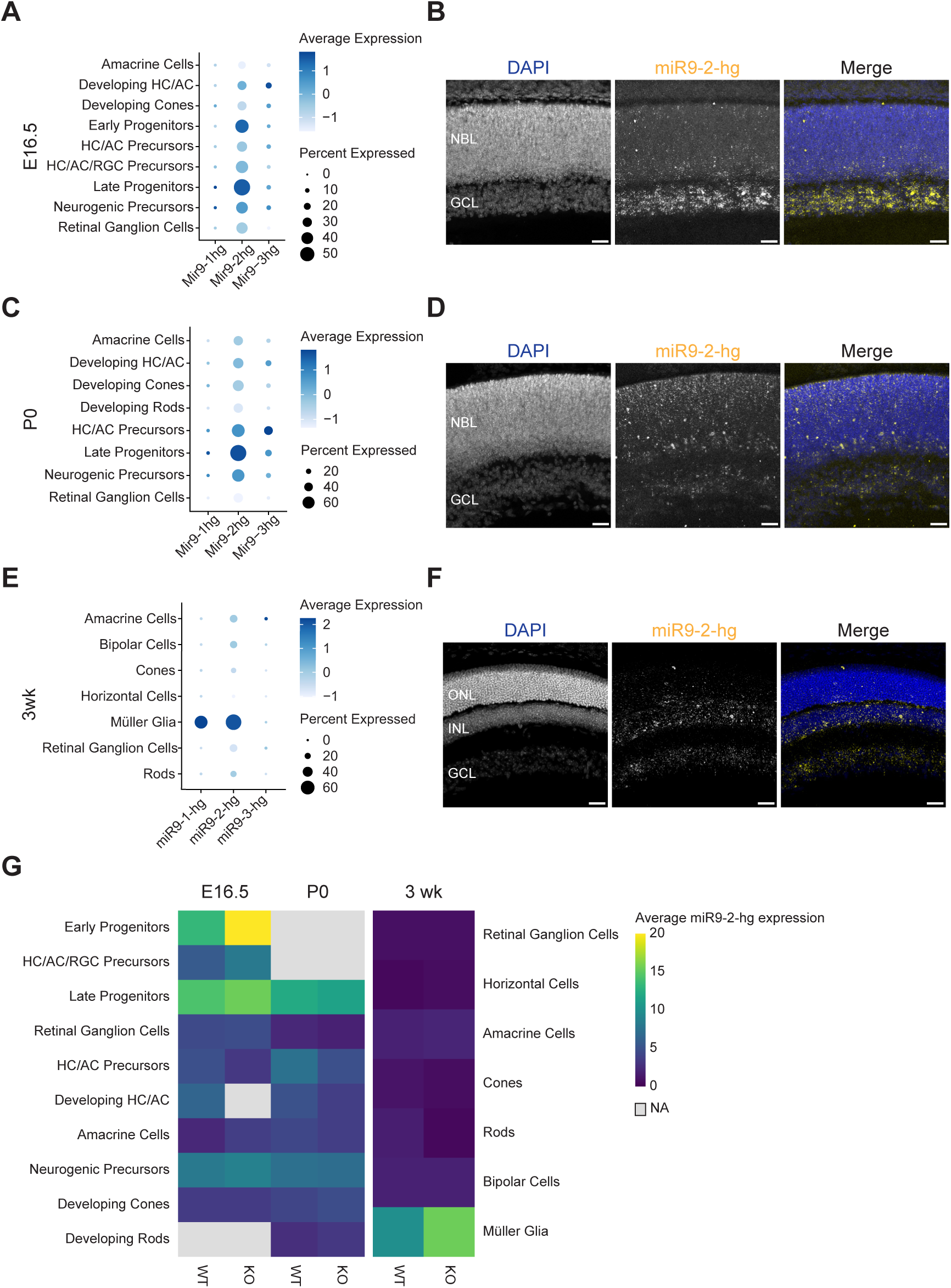
***Mir9-2hg* is enriched in retinal progenitor cells and Müller glia.** Dot plots of miR-9 paralog expression in WT cells of major cell classes from snRNA-seq at E16.5 **(A)**, P0 **(C)**, and 3-weeks **(E)**. *In situ* RNA hybridization images of *Mir9-2hg* (yellow) in cross sections of WT C57Bl6/J animals at E16.5 **(B)**, P0 **(D)**, and 3-weeks **(F)**. Scale bars = 50 microns. **G.** Single-nucleus RNA-seq heatmap of average expression of *Mir9-2hg* in WT and miR-9-2 KO mice in each cell class. Gray boxes indicate cell classes in negligible numbers/not present. Scaled (0-20) across all timepoints. Abbreviations: HC = horizontal cells, AC = amacrine cells, RGC = retinal ganglion cells, NBL = neuroblastic layer, GCL = ganglion cell layer, ONL = outer nuclear layer, INL = inner nuclear layer.

We aimed to validate the miR-9-2 expression patterns using fluorescent *in situ* hybridization. We collected wildtype C57Bl6/J tissue at timepoints aligning with our snRNA-seq data and used RNAscope Plus to probe for *Mir9-2hg* transcripts. At E16.5 and P0, the expression of the transcript was dispersed across the neuroblastic layer of the retina (Fig. 3B, 3D). The positioning of these transcripts is reflective of the migratory movement of RPCs during development as they migrate apically and basally during division and proliferation^44,45^. Additionally, at E16.5 there appeared to be strong labeling in the developing ganglion cell layer, potentially corresponding to cytoplasmic accumulation in post-mitotic precursor cells (Fig. 3B). At 3-weeks, the expression of the transcript became more concentrated in the inner nuclear layer (Fig. 3E), consistent with the MG-specific expression seen by snRNA-seq and lacZ reporter mouse expression (Fig. 1E).

We next sought to understand how this expression was affected in the miR-9-2 KO and whether other miR-9 paralogs would be able compensate for loss of miR-9-2. Using our snRNA-seq dataset, we found that *Mir9-2hg* remained the predominant paralog expressed in the mouse retina across all time points in WT and KO animals (Fig. 3G; SFig. 7). Unexpectedly, expression of *Mir9-2hg* appeared to be increased in KO retinas, particularly in early progenitors at E16.5 and MG in the mature retina (Fig. 3G). This implies a potential autoregulatory role of miR-9, where disruption of an autoinhibitory signal from mature miR-9 causes an increase in *Mir9-2hg* expression. Such a negative feedback loop for miR-9 has been suggested in other systems but not directly demonstrated in mammals.

### Loss of miR-9-2 leads to direct and indirect target gene disruption during the transition from early-to-late born retinal cell classes

After identifying the major cell types that express miR-9-2, we sought to characterize how each developmental period was affected by miR-9-2 loss. To do so, we performed differential expression analyses using Presto^46^ to compare WT and miR-9-2 KO transcriptional profiles of each cell class (Table S3). As the primary function of a miRNA is inhibiting translation of mRNAs through degradation or translational block of target transcripts with perfect or imperfect complementarity^21^, we expected that loss of a miR-9-2 would disrupt a large set of putative target genes. To distinguish between direct and indirect targets of miR-9-2, we first assembled computationally predicted targets of miR-9 (both 3p and 5p targets) from the TargetScanMouse 8.0 database^47,48^. Next, we determined which of these predicted targets were expressed in WT and/or miR-9-2 KO cells. For direct targets, we expected that loss of the miRNA would lead to de-repression and thereby upregulation of target genes in miR-9-2 KO cells compared to WT. Concordant with these expectations, we found de-repression of specific target genes and widespread dysregulation of other indirect targets across timepoints and cell classes.

At E16.5, miR-9-2 expression begins to rapidly increase during mouse retinal development^22^, especially within RPCs. In early RPCs, some of the top upregulated genes included the TFs *Rorb* and *Meis1* (Figure 4B) which are predicted to be direct miR-9 targets. Furthermore, late RPCs have significant upregulation of additional putative miR-9 target TFs including *Foxp1* and *Pbx1*, and early upregulation of Onecut family of TFs (*Onecut1/2/3*; not shown, p≤0.05; Figure 4C). Many of these TFs already have well documented roles in regulating neurogenesis, in CNS and in retina specifically. Outside of TFs, some of predicted miR-9 targets that were highly upregulated include *Auts2* and *Ank2,* which is consistent across early and late RPCs. These genes have been implicated in neurodevelopment and have documented roles in neuronal migration and synaptic formation and function^49,50^.

**Figure 4.**
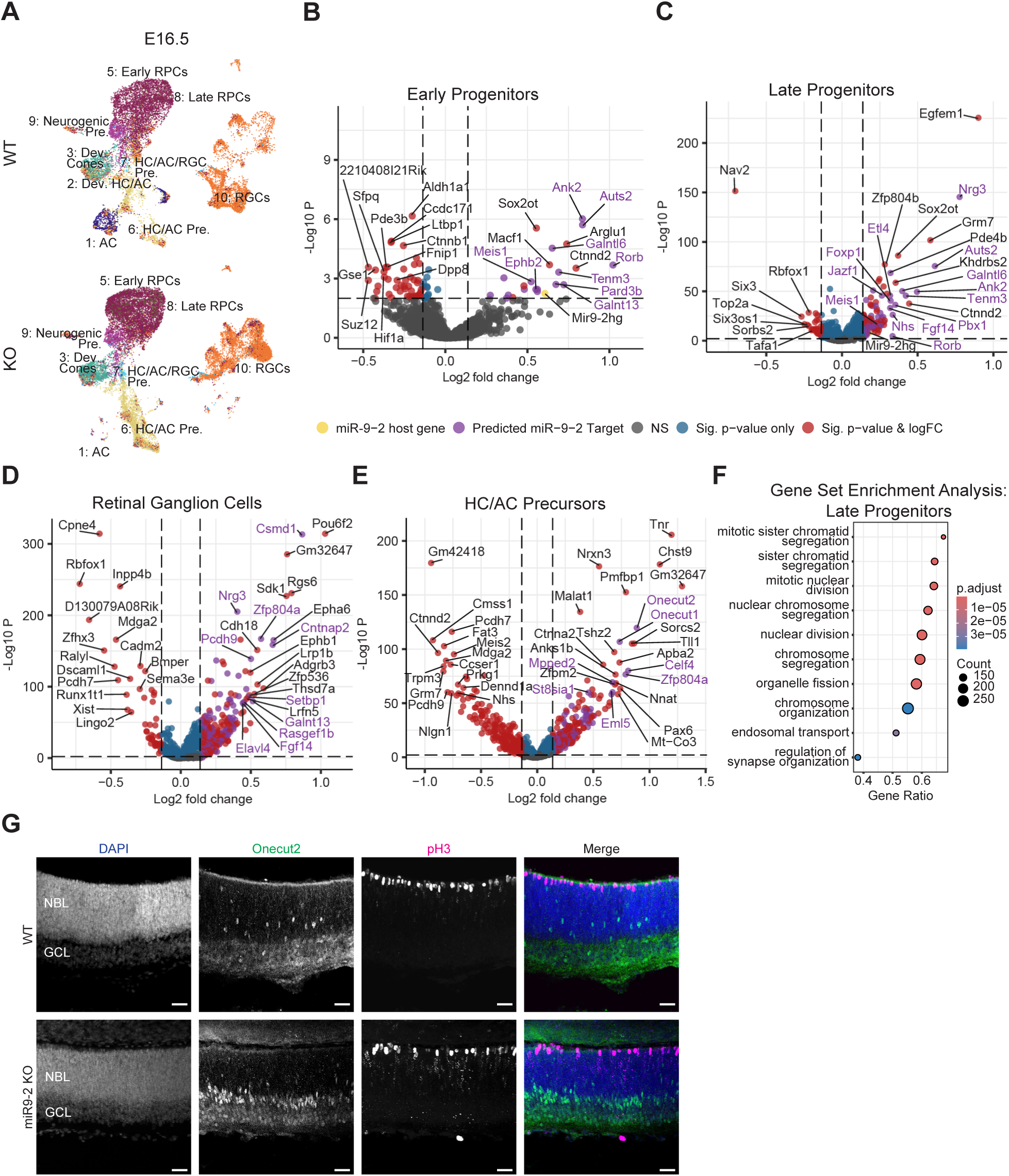
Transcriptional profiles of cell classes in miR-9-2 KO retinas are disrupted at E16.5. **A.** UMAP of cell classes in miR-9-2 KO and WT retinas at E16.5. **B-E.** Volcano plots of differentially expressed genes in individual cell classes at E16.5 including early RPCs **(B)**, late RPCs **(C)**, RGCs **(D)**, and HC/AC Precursors **(E)**. Thresholds (dashed lines) correspond to p≤0.01 and FC≥10%. Wilcoxon rank sum test, Benjamini-Hochberg correction. **F.** GSEA of the top 10 enriched terms in E16.5 late RPCs. Adaptive multi-level split Monte-Carlo^92^. **G.** Immunolabeling of ONECUT2+ (green) and pH3+ (magenta) cells of E16.5 KO and WT retinas in cross-section. All scale bars = 50 microns. Abbreviations: RPC = retinal progenitor cells, RGC = retinal ganglion cells, HC = horizontal cells, AC = amacrine cells, Pre. = precursor, Dev. = developing, NBL = neuroblastic layer, GCL = ganglion cell layer.

Since loss of miR-9-2 has wide-ranging effects beyond direct target de-repression, rather than focusing on individual genes, we performed pathway analyses for each cell class to give a broader sense of which processes are affected in miR-9-2 KO retinas. We performed a Gene Set Enrichment Analysis (GSEA) using the ClusterProfiler package^51^ with the Gene Ontology Biological Processes (GOBP) database^52,53^ of pathways for reference. Of the top 10 enriched pathways for late RPCs, most terms were associated with mitosis including “mitotic sister chromatid segregation” and “mitotic division” (Fig. 4F). This enrichment of mitotic pathway terms in KO retinas could suggest increased proliferation, delay of cell cycle exit, or a difference in cell cycle length which would be in alignment with a more immature state. Using pH3 as a marker of dividing cells, we immunolabeled developing E16.5 retinas and found that KO retinas appeared to have a sparser distribution of dividing cells on the apical (choroidal) side of the neuroblastic layer, but an increased number of cells containing condensing chromatin dispersed throughout the neuroblastic layer (puncta; Fig. 4G). To investigate further, using Seurat, we performed cell cycle phase analysis, which classifies cells into G1, G2/M, or S phases, and found only one cell class, Developing Cones, that differed in G2M phase proportions between WT and KO retinas (SFig. 5; Table S2). However, for cones, especially as miR-9-2 expressing RPCs aren’t affected, it seems likely that this result could be related to the overall alteration in cell class production rather than direct cell cycle modulation. These data suggest, therefore, that miR-9-2 could be controlling cell class emergence independently of cell cycling states.

While changes to RPCs are clear, we also wanted to understand how these alterations affect production of retinal cells following specification of retinal cell types. As *Mir9-2hg* expression is highest in RPCs and upregulated genes in early RPCs are significantly enriched for computationally predicted miR-9-2 targets (Table S6), we expect later cell type alterations to be the result of upstream progenitor changes. Retinal ganglion cells (RGCs) are the earliest born cell type of the retina, and by E16.5, the majority of RGCs have been born. Between miR-9-2 KO and WT retinas, there is a clear segregation of RGC populations visible on the E16.5 UMAPs (Fig. 4A). To understand why these populations are distinct, we compared the transcriptional profiles of KO and WT RGCs (Fig. 4D). The top upregulated gene in KO RGCs is *Pou6f2*, a ganglion cell marker that is expressed in immature RGCs and ACs during development and only develops a restricted expression pattern to RGC subtypes in adult retinas^54,55^. Within individual cell classes, cell types are also born with a distinct order and timing; therefore, these data suggest that not only is the overall retina more immature, but also that individual cell types are immature as well.

As the AC population is decreased in E16.5 miR-9-2 KO retinas, we focused on horizontal and amacrine precursor cells (HC/AC Precursors) to understand what leads to this alteration. Many of the top upregulated genes in these precursors persist from early and late RPCs and include the TFs, *Onecut1* and *Onecut2* (Fig. 4E). Onecut TFs are among the top computationally predicted targets of miR-9 and have been experimentally confirmed to be direct targets of miR-9 in the CNS^29,47,48^. To validate this finding, we immunolabeled E16.5 retinas for ONECUT2 and found a significant increase in ONECUT2+ cells in KO retinas compared to WT (Fig. 4G). In the context of retinal cell fate specification, Onecut TFs have well-characterized roles in specifying early-born cell classes, including ACs^8,9,13^. Altogether, these data support the hypothesis that mir9-2 KO causes a developmental delay in retinal cell class specification and differentiation.

### Loss of miR-9-2 leads to dysregulation of late-born cell classes in the developing mouse retina

To see how the development of the retina is affected during the peak of late-born cell birth, we profiled WT and miR-9-2 KO retinas at P0. By this stage, RPCs are primarily giving rise to post-mitotic neurons, such as rods, bipolars, and late-born amacrines, and later, MG^1,4^. This timepoint is also during a plateau of high *Mir9-2hg* expression in the developing retina. We again compared the transcriptional profiles of WT and KO cells with a focus on RPCs and precursors that are implicated in the production of late-born cell types (Table S4). While several of the genes upregulated in E16.5 late RPCs remain persistently upregulated, like *Meis1*, *Ank2*, and *Auts2*, at P0, *Nfib* becomes one of the top downregulated genes in late RPCs. While more moderate, there is also a significant downregulation of other NFI factors, *Nfia* and *Nfix* (Fig. 5B). NFI factors, while not predicted to be direct miR-9 targets, are found to have late RPC-specific expression and have been implicated in specification of late-born cell classes, such as BCs and MG^19^. Additionally, other genes with documented roles in regulating neural progenitor proliferation and differentiation, including *Notch1*, *Hes1*, *Rbpj*, and *Notch2* (*Rbpj*, *Notch2*; not shown, p≤0.05)^14–16,18^, are also significantly decreased in P0 late RPCs (Fig. 5B). Disruptions of downstream gene networks regulated by these factors likely contributes to further alterations in RPCs and the post-mitotic neurons they produce.

**Figure 5.**
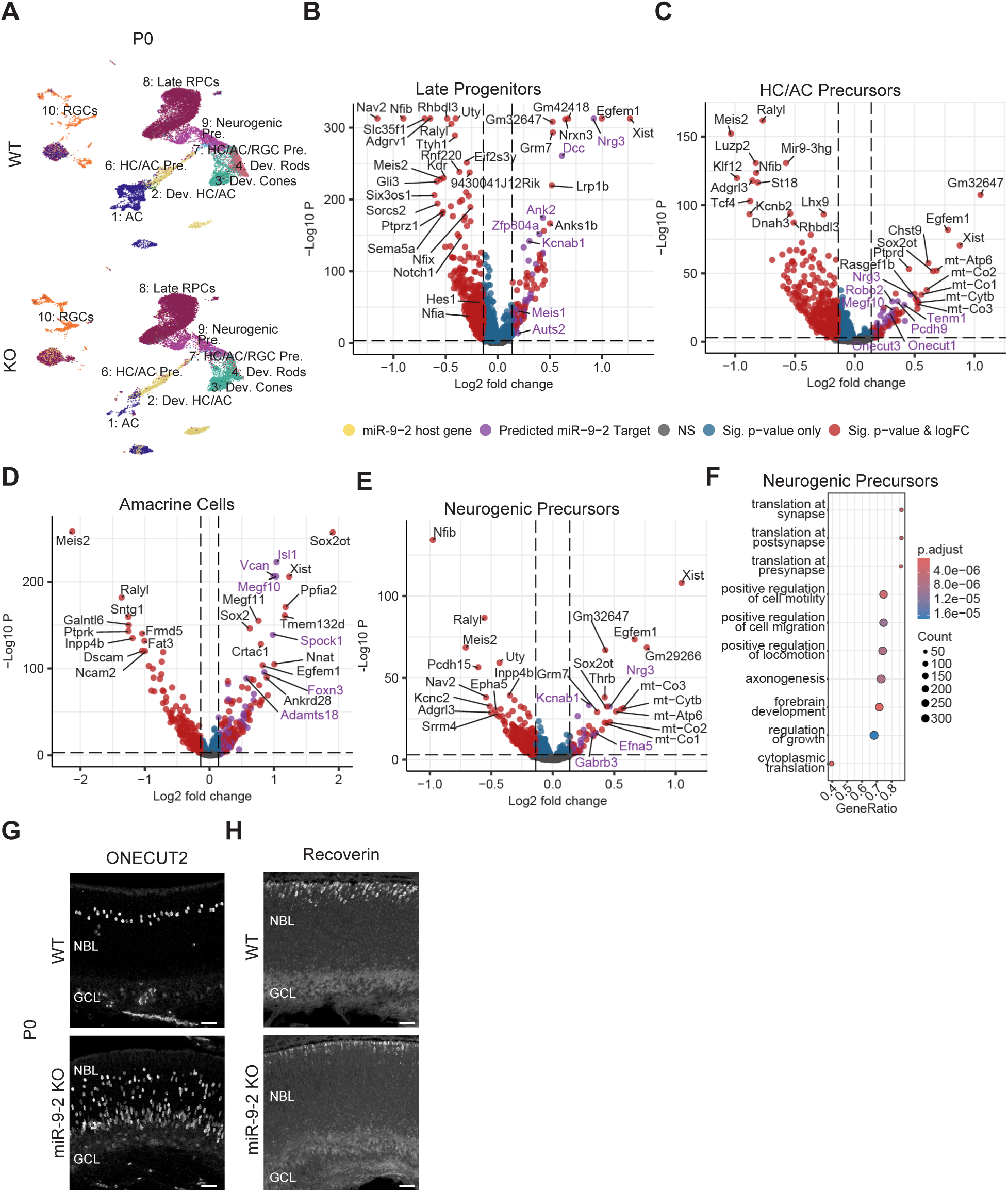
Transcriptional profiles of cell classes in miR-9-2 KO retinas are disrupted at P0. **A.** UMAPs of cell classes in P0 miR-9-2 KO and WT retinas. **B-E.** Volcano plots of differentially expressed genes in individual cell classes at P0 including late RPCs **(B)**, HC/AC precursors **(C)**, ACs **(D)**, and neurogenic precursors **(E)**. Thresholds (dashed lines) correspond to p≤0.01 and FC≥10%. Wilcoxon rank sum test, Benjamini-Hochberg correction. **F.** GSEA of the top 10 enriched terms in neurogenic precursors. Adaptive multi-level split Monte-Carlo^92^. **G.** Immunolabeling of ONECUT2+ cells in P0 KO and WT retinas (WT: n=5; KO: n=6) in cross-section. Mean +/- 1SD. Wilcoxon rank sum text, **p≤0.01. **H.** Immunolabeling of Recoverin+ cells (green) in P0 KO and WT retinas. All scale bars = 50 microns. Abbreviations: RPC = retinal progenitor cells, RGC = retinal ganglion cells, HC = horizontal cells, AC = amacrine cells, Pre. = precursor, Dev. = developing.

Among post-mitotic cells, developing HC/ACs and distinct ACs emerged by P0 in both WT and KO retinas (Fig. 5A). However, knockout ACs and their precursors still appear biased towards earlier born HC/AC types compared to their WT counterparts. In HC/AC precursors, as at E16.5, the Onecut TFs are still significantly upregulated (Fig. 5C, 5G). In KO retinas, we saw a significant increase in ONECUT2+ cells compared to WT. Further, while ONECUT2+ cells in WT retinas are organized in two distinct layers corresponding to HCs and ACs, in KO retinas, ONECUT2+ cells are more widely distributed. This disorganized state is characteristic of immature cells, as RPCs and precursors migrate apically and basally until they are fully matured. The ACs in KO retinas also have increased expression of early-born AC markers, and the top upregulated genes include *Isl1*, *Sox2*, *Megf10*, and *Megf11* which are specifically expressed in cholinergic starburst ACs, an early-born AC type^56–59^. Like the more immature RGCs at E16.5, these data indicate that ACs in KO retinas are dominated by earlier AC types.

Outside of these progenitor and amacrine cell alterations, the most striking effect at P0 is the near absence of developing rods in miR-9-2 KO retinas (Fig. 5A). To elucidate the cause, we compared neurogenic precursor cells, a population that can adopt photoreceptor fate at P0, between WT and KO retinas. *Thrb*, a marker of cones, is one of the top upregulated genes in KO neurogenic precursor cells, suggesting that these cells are more primed to adopt a cone fate, which is an earlier-born photoreceptor cell class compared to rods (Fig. 5E). When we performed a GSEA on these precursors, we found that some of the top enriched terms included “translation at the synapse”, “positive regulation of cell motility and migration”, and “axonogenesis” (Fig. 5F). These terms are indicative of processes related to neuronal maturation, as differentiating neurons need to migrate to the correct spatial location, locate synaptic partners and form functional circuits. Lastly, to validate changes to photoreceptors in KO retinas, we performed immunolabeling for recoverin, a marker of retinal photoreceptors (Fig. 5H). In alignment with our snRNA-seq data, we find a decrease in the number of recoverin-positive cells in the KO compared to WT. This supports the observation that loss of miR-9-2 delays formation of rods. Altogether, these data collected at the peak of late neurogenesis suggest that miR-9 is important for both controlling cell class emergence and also controlling the timing of cell class subtype specification.

### Loss of miR-9-2 leads to dysregulation of MG cells in the mature mouse retina

Despite the striking developmental delay in miR-9-2 KO developing retinas, by 3-weeks, the proportions of major cell classes are not significantly different (Fig. 2, Fig. 6A). Paradoxically, this result appears to be in direct conflict with our KO retinal histology, where we find an increase in Sox9+ cells suggesting an increase in MG. To resolve this incongruency, we first compared the transcriptional profiles of WT and KO Müller glia (Fig. 6B; Table S5). Among the top upregulated genes, we found several predicted miR-9 targets including *Cp*, *Trpm3, Foxp2*, and *Dxd5,* among other genes including *Apoe*, *Nrbp2*, *Cadbps2*, and *Lrmda*. Top downregulated genes included *Prkcb*, *Rora*, *Nrxn3*, and *Spc25*. We performed a GSEA, to understand what processes were disrupted in these cells. Some of the top enriched terms included “translation at synapse”, “cellular respiration”, and “mononuclear cell differentiation” (Fig. 6C). Disruptions to these pathways could suggest a disruption to Müller identity, where enrichment of synapse formation and changes to metabolic processes could suggest that KO Müllers have more neuronal attributes. We also performed a TF enrichment analysis using CHEA3^60^ and found that among the top predicted TFs, there are putative miR-9 targets, including *Satb2* and *Rest*, which have documented roles in chromatin remodeling^61,62^ ^63^ (Fig. 6D). Taken together, these data provide further evidence for the altered state of KO MG and is in alignment with our early findings that KO MG co-express both Müller markers and neuronal (BC) markers (Fig. 1E-F).

**Figure 6.**
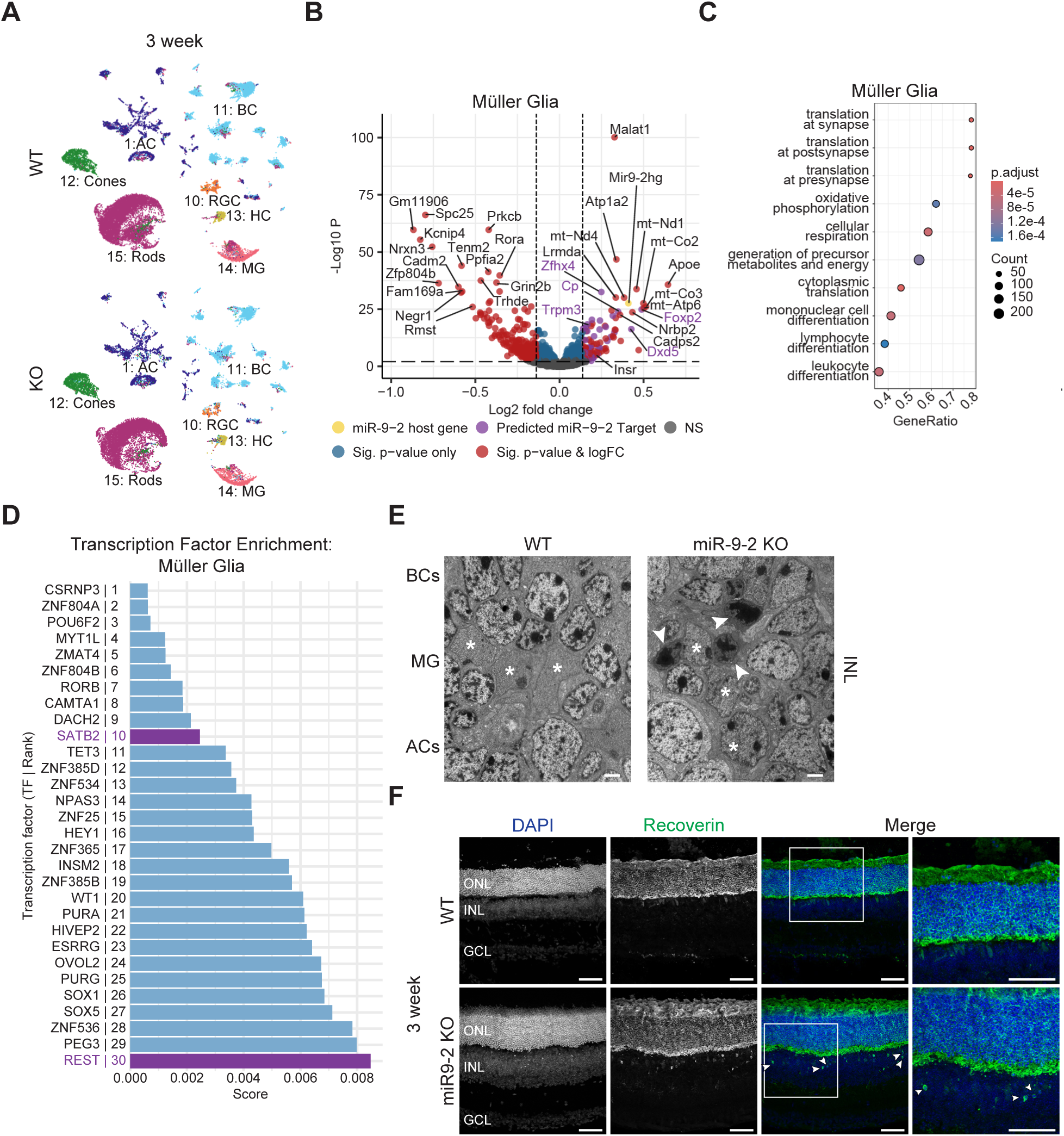
3-week mature MG cells are disrupted in miR-9-2 KO retinas. **A.** UMAPs of cell classes in 3-week miR-9-2 KO and WT retinas. **B.** Volcano plot of differentially expressed genes in 3-week MG. Thresholds (dashed lines) correspond to p≤0.01 and FC≥10%. **C.** GSEA of the top 10 enriched terms in 3-week MG. **D.** Top 30 ranked genes from TF enrichment analysis (rCHEA3) for MG. Low-scoring TFs contribute most. Over-representation analysis, Fisher’s Exact Test. TFs that are upregulated in KO and are computationally predicted miR9 targets in purple. **E**. Transmission electron microscopy images of INL of WT and KO retinas. MG cells (white asterisks), ACs, and BCs are visible. Rods indicated by white arrowheads. Scale bars = 2 microns. **F**. Immunolabelling of Recoverin+ (green) photoreceptors in KO and WT retinas in cross-section. Rods in INL marked by white arrowheads. Lower expressing recoverin+ BCs are seen in both WT and KO. Scale bars = 50 microns. Abbreviations: AC = amacrine cells, BC = bipolar cells, RGC = retinal ganglion cells, HC = horizontal cells, MG = Müller glia, ONL = outer nuclear layer, INL = inner nuclear layer, GCL = ganglion cell layer.

To further probe the alterations to 3-week retinas, we used transmission electron microscopy (TEM) to investigate MG morphology. We harvested mature retinas from 3-week-old mice and assessed for gross morphological differences between WT and miR-9-2 KO animals. Focusing on MG, we found that the nuclei of KO cells appeared altered. MG nuclei are characterized by their polygonal shape and linear positioning in the middle of the INL, with their heterochromatin normally appearing uniformly diffuse across the nucleus (Fig. 6E, asterisks, WT). However, in KO retinas, MG chromatin looks like the surrounding interneuron nuclei (ACs, BCs), where chromatin has multiple focally condensed chromocenters within each nucleus (Fig. 6E, asterisks, KO). KO retinas also appeared disorganized, with infiltration of rods into the INL even though these nuclei would normally only reside in the ONL. Rod nuclei have a distinctive ring-like appearance as rod heterochromatin is condensed to a large central mass^64^ (Fig. 6E, arrowheads). To further validate our findings, we performed immunolabelling for markers of rods in 3-week KO and WT animals. As in the TEM, we found recoverin+ cells abnormally localized in the INL (Figure 6F, arrowheads). Like our snRNA-seq data, these data indicate a disorganization of normal retinal structure and point to potential alterations in cell class identity, especially as mature MG cells no longer display their characteristic chromatin patterning, instead taking on a more neuronal appearance. Further, as MG often play a structural support role in the retina, this disruption to the glia may contribute to rod mis-localization and disruption of retinal architecture. Dysregulation of glial identity and disruption of critical pathways such as oxidative phosphorylation, cellular respiration, along with de-repression of neuronal programs in miR-9-2 KO MG may shed light on mechanisms of retinal disease risk associated with miR-9-2 and its conserved enhancer.

## DISCUSSION

The importance of microRNAs in generating a fully functional retina has been well established, but it is not fully known which miRNAs contribute most to developmental regulation and which aspects of development they regulate. In this work, we developed an *in vivo* germline knockout mouse model of miR-9-2 function and found that, disruption of miR-9-2 delays the timing in which retinal cell classes emerge. Surprisingly, we find this delay occurs not just for the major cell classes, but also for cell types within those classes, such as cholinergic and glycinergic ACs. Additionally, in the absence of miR-9-2, we determined that, while all retinal cell classes will eventually form, the MG of miR-9-2 KO animals appear to be mis-specified and adopt a hybrid neuronal- and Müller-like cell fate. This suggests that miR-9-2 sits at a critical intersection of GRNs to control retinal progenitor competence with a high degree of temporal precision.

We identified several key TFs that miR-9-2 directly regulates to control these cell fates, including the Onecut, Rorb, and Foxp families. Of these key regulators, all are computationally predicted to be direct target genes of miR-9-2 and are de-repressed in miR-9-2 KO RPCs, the cell type with highest miR-9-2 expression in development. Several putative miR-9-2 targets included TFs that are known to promote progenitor proliferation, such as *Rorb*, or TFs that promote early-born cell fates, like *Foxp1* and the Onecut TFs (including *Onecut1/2/3*) ^8–10,12,13,65,66^. In addition to interneurons, Onecut TFs and *Rorb* have documented roles in regulating photoreceptor fates, including promotion of cone cell fate (former) and maintenance of the balance between cone and rod fates (latter)^8,12,13,67,68^. With the loss of miR-9-2, these TFs were de-repressed, likely contributing to the increased proportions of early-born cell classes and the overall immature phenotype of the KO retinas. Therefore, the direct interactions of miR-9-2 with these crucial TFs seem to contribute to the ability of RPCs to transition from producing early- to late-born cell fates at the proper developmental time.

More broadly, the loss of miR-9-2 appeared to target genes and pathways that are found to be highly involved in retinal cell fate specification. Specifically, expression of Notch signaling effectors was downregulated in miR-9-2 KO late RPCs, including a large (25%) decrease in *Notch1* with more subtle, yet still significant decreases in expression of *Hes1*, *Notch2,* and *Rbpj* (Fig. 5B). Notch expression can promote continued proliferation of RPCs and regulate the formation of early-born cell types like RGCs and cones^14–16,18,28,69^. Late RPCs also had a substantial downregulation of NFI factors. This may be driven, in part, by the upregulation of the TF, *Foxp1*, in E16.5 late RPCs (Fig. 5B), as its overexpression has been found to inhibit *Nfib*^70^. Recent single cell studies found that NFI factors are restricted to late RPCs during retinal development and may promote MG and BC fates^19^. Therefore, in addition to directly regulating key drivers of early-born cell fate, miR-9-2 seems to indirectly regulate the proper balance of TFs and signaling pathways that are important for promoting late-born fate.

By the end of development, while all retinal cell classes eventually formed, the presence of mis-specified MG cells in the mature miR-9-2 KO retina was distinct. We found mis-expression of OTX2, choroidal migration of SOX9+ MG to the ONL, and qualitative changes in the chromatin structure of these cells. Altogether, these data suggest that KO MG have a neuronal-glial hybrid identity and seem to be developing a more neuronal fate. The hybrid state of MG may also contribute to our inability to see cell proportion changes in our snRNA-seq datasets as it is probable that hybrid cells segregate into MG and BC clusters. Transcriptionally, of the DEGs identified in MG of miR-9-2 KO and miR-9-2 enhancer knockout mice, there are only four upregulated putative miR-9 targets shared between models, including *Trpm3* (addressed below), *Foxp2*, *Insr*, and *Cdh4*. *Insr*, *Cdh4*, and *Foxp2* have all been implicated in synapse formation in the CNS^71–74^ while *Foxp2*, in particular, is a TF that has also been linked to speech and motor circuit formation and neurogenesis^75^. In regenerative animals such as zebrafish, MG act as latent RPCs, and in response to injury can de-differentiate to become neurons. While mammalian MG do not retain this native ability, they are known to have progenitor-like expression profiles and have some neurogenic potential when artificially induced to differentiate. Therefore, miR-9-2 seems to have a role in maintaining the progenitor-like identity of these MG, and its loss may be permissive for differentiation into other cell fates. The MG-restricted expression of miR-9-2 in the mature retina suggests that miR-9-2 may become important for the maintenance and homeostasis of these cells. Further work is required to explore the potential of miR-9-2-related modifications to MG identity and determine whether miR-9 depletion can be used regeneratively, to promote Müller glia reprogramming. As this paper models the disruption of miR-9-2 in all tissues, from conception, it is possible that more targeted disruptions to miR-9 function, occurring at later timepoints or targeting specific cell types (such as Müller glia) could provide great value in future study.

Lastly, this mouse model of miR-9-2 loss serves as a starting point for understanding retinal disease risk. The miR-9-2 locus has been increasingly linked to Macular Telangiectasia Type II (MacTel), a rare age-related degenerative disease that leads to central vision loss. In MacTel, early-stage patients appear to lose MG cells prior to photoreceptor and subsequent vision loss while also experiencing metabolic defects^34,76,77^. In this study, we found several genes upregulated in miR-9-2 KO MG that could contribute to this disease, including *Cp*, a copper transporter and iron oxidoreductase^78^, and *Trpm3*, a Ca2+ channel that can be activated by sphingosine^79^. In the context of MacTel, dysregulation of *Cp*, and thereby iron homeostasis, could lead to inflammation and/or ferroptosis^78^, while *Trpm3* over-activation by sphingolipids could cause eventual loss of MG in MacTel patients^77^. *Trpm3* is also the host gene of another miRNA, miR-204. As *Trpm3* is upregulated in KO retinas, we looked for repression of its predicted targets and found that significantly downregulated genes in KO MG were enriched for predicted miR-204 targets (Table S6). While this was an unexpected finding, it solidifies the concept that miR-9-2 sits at a vital intersection of GRNs which allows it to have these wide-ranging effects. This study highlights how a single individual miRNA can have a significant impact on proper retinal development and its risk for retinal disease.

### Limitations of the study

In this study, we find new insight into the function of an individual miRNA, miR-9-2, and its effects on retinal development and glial identity. However, whole body disruption of miR-9-2 led to additional unexpected phenotypes in brain and affected animal survival (see Fregoso et al.) which made it difficult to capture much of the postnatal development of these mice and led to small sample size. We also were only able to study retinas from development to early maturity, even though many retinal diseases are late onset. Similarly, this made it difficult for us to parse if miR-9-2 has distinct roles in development versus glial homeostasis. Future studies would benefit from targeted disruptions of miR-9-2 function and investigation into miR-9-2 function in aging retinas.

## METHODS

### Mice

#### Generation of miR-9-2 knockout animals

*MiR-9-2^tm1Mtm^* mice, containing a beta-galactosidase reporter for miR-9-2, a neomycin selection cassette, and the floxed miR-9-2 hairpin loop were obtained from Jackson Laboratories^41^. To remove the selection cassette and beta-gal reporter which are flanked by Frt sites, we first crossed *MiR-9-2^tm1Mtm^* mice with *ROSA26::FLPe* knock in mice^42^; subsequently, to knockout miR-9-2, we crossed the resulting progeny with *EIIa-cre* mice^1^. This resulted in a germline miR-9-2 knockout (KO) strain with constitutive loss of miR-9-2 in all cells (see Table S7 for genotyping primers for all lines). To ensure uniform C57Bl6 background, heterozygous animals were backcrossed to C57Bl6 breeders for at least 10 generations.

#### Sample harvest

To obtain retinas from embryonic timepoints, breeding animals were paired and plug checks performed daily. Once plugs were confirmed, females were separated from males and housed individually. Females were monitored to confirm pregnancy before embryos were extracted at embryonic age (E), E16.5. To obtain postnatal timepoints, breeders were monitored for birth daily, and pups were collected at relevant postnatal (P) timepoints, P0 and 3-weeks (P21-P25). For immunofluorescence assays, P0 and E16.5 whole heads were harvested, or 3-week eyes were enucleated and immediately fixed post-harvest. For all single cell samples, eyes were enucleated, retinas dissected out, and tissue immediately flash frozen in liquid nitrogen and stored at −80C or in liquid nitrogen until use. For RNAscope samples, we used C57Bl6/J timed pregnant dams from JAX (JAX:000664) to harvest wildtype retinas at E16.5, P0, and 3-week timepoints. Retinas from these animals were collected and harvested for immunohistochemistry as described above.

### Immunohistochemistry

#### Fixation and cryosectioning

<u>Fixation</u>: For E16.5 and P0 whole head samples, tissue was fixed for 24 hours at 4C. For 3-week samples, enucleated mouse eyes were fixed for 1 hour at room temperature with cornea opened to expose tissues. All samples were fixed in a solution of 4% PFA in 1X PBS. After fixing, samples were washed 3 times with 1X PBS. <u>Cryosectioning:</u> For enucleated eyes, cornea and lens were dissected off. Retinas and whole heads were then washed through a sucrose gradient of 5%, 15%, and 30% sucrose in 1X PBS until tissues sank in solution. Tissue was embedded in 100% O.C.T., frozen on dry ice, and stored at −80C. Cryosectioning of 40-micron sections was performed at −20C, and sections were mounted on TruBond Gold slides, dried on a 37C slide warmer for 1 hour, and stored at −20 to −80C until use.

#### Antibody staining

Upon removal from storage, before staining, slides were dried for 3 hours at 60C to maximize tissue adherence. To remove O.C.T., tissue was rinsed for 15 minutes in 1X PBS. To promote antigen retrieval, slides were steamed for 10 minutes in sodium citrate buffer (10mM Citric Acid, 0.05% Tween 20, pH 6.0) and washed with 1X PBS to prepare for antibodies. For primary antibody (see table for antibodies and dilutions), antibodies were diluted into blocking solution (1% BSA, 0.05% SDS, 0.01% Triton in 1X PBS) and left on slides at 4C overnight in a humidified chamber. Primary antibody was removed by washing slides 3 times with 1X PBS. Secondary antibody was diluted (1:750) in blocking solution and left on tissue for 2-3 hours at room temperature (RT). Antibody was removed by washing 3times with 1X PBS and slides were cover-slipped using Prolong Gold Antifade.

#### TUNEL staining for cell death

TUNEL assays were performed using the *In situ* Death Detection Kit, Fluorescein (Roche). Experimental slides prepped for immunohistochemistry along with two control (positive and negative) slides were used for each experiment. The positive control tissue was treated with DNase I recombinant, grade I at a concentration of 300U/mL in 50mM Tris-HCl, pH 7.5, 10mM MgCl2, 1mg/mL BSA for 10 minutes at 37°C to induce DNA strand breaks prior to labeling. All slides were washed 30 minutes with 1X PBS then permeabilizated in 0.2% Triton X-100 in fresh 0.1% sodium citrate solution at RT for 30 minutes. Slides were rinsed twice with 1X PBS and then excess liquid removed. In a humidified chamber, 150uL of TUNEL reaction mixture (1:9 ratio of Enzyme to Label solution) was applied to each slide. The negative control was treated only with the Label solution. Slides were then incubated for 60 minutes at 37°C, washed 3 times with 1X PBS, then stained with DAPI antibody at a 1:1000 concentration for 10 minutes. After a 1X PBS wash, slides were cover-slipped using Prolong Gold antifade.

### *In Situ* Hybridization

*In situ* hybridization was performed using RNAscope Plus smRNA-RNA HD Reagent Kit (ACD). Tissue was fixed as described previously, cryosectioned into 20-micron sections at −20°C, and then mounted on Trubond Gold slides. Slides were air-dried for 60 minutes at RT, then washed for 5 minutes with 1X PBS before baking for 60 minutes at 60°C in a hybridization oven. Baked slides then underwent post-fixation in 4% PFA in 1X PBS for 15 minutes at 4°C. To dehydrate tissue, slides were put through an ethanol gradient (50%, 70%, 100% EtOH) for 5 minutes each at RT before air-drying for 5 minutes at RT. Tissue was post-fixed in 4% PFA in 1X PBS overnight at 4°C. Slides were then treated with RNAscope Hydrogen Peroxide for 10 minutes at RT and washed twice in distilled water. To perform target retrieval, slides were first immersed in prewarmed distilled water for 10 seconds to acclimate to temperature before being immersed in prewarmed 1X Target Retrieval Reagent and steamed for 10 minutes. They were promptly removed and rinsed in distilled water for 15 seconds, immersed in 100% EtOH for 3 minutes, then air-dried at RT for 5 minutes. RNAscope Protease III was applied to each slide for 10 min at 40°C in a hybridization oven, then rinsed twice with distilled water. ACD performed probe design with the S1 probe targeting the miR-9-2 host gene. Slides were covered in the probe solution, allowed to hybridize for 2 hours at 40°C, then rinsed twice in 1X Wash Buffer. For amplification, RNAscope Plus smRMA-RNA HD AMP1 was added to slides for 30 minutes at 40°C, then washed twice in 1X Wash Buffer. This step was repeated for AMP2 then AMP3, however AMP3 was only incubated for 15 minutes. To develop the signal, RNAscope Plus smRMA-RNA HRP-S1 was applied for 15 minutes at 40°C, and washed twice in 1X Wash Buffer. To conjugate the fluorophore to probe, the slide was incubated with TSA Vivid Fluorophore 520 dye (1:500, in TSA buffer) for 30 minutes at 40C. Slides were rinsed twice in 1X Wash buffer, incubated in HRP Blocker for 15 minutes at 40°C, and rinsed twice again with 1X Wash Buffer. After the final rinse, DAPI was added to each slide for 30 seconds at RT then cover-slipped using Prolong Gold antifade. Coverslips were dried at RT overnight and protected from light before imaging or storage at −20°C.

### Electron Microscopy

Eyes were enucleated from 3-week miR-9-2 knockout and wildtype littermate pairs and fixed in 4% glutaraldehyde in 0.1M cacodylate and then washed 4 times in 0.1M sodium cacodylate buffer. Samples were post-fixed in 2% osmium tetroxide, buffered, followed by 1% uranyl acetate in water, and then washed 4 times with water. Tissue was dehydrated in a series of ethanol washes from 30-100% ethanol and then given two changes of propylene oxide. Tissue was embedded in Epon Araldite 502 and cured for 24 hours at 60C. A JEOL JEM 1200EXII electron microscope at 80kV and a spot size of three was used to image 80nm sections.

### Confocal Microscopy and Image Processing

#### Microscopy

All slides were imaged using the Leica Sp5 or Zeiss LSM 900 or 980 confocal microscopes. Images were taken using a 20X air or 40X oil immersion lenses, with Z stacks taken at 2-micron increments, spanning the entire thickness of the section/retina.

#### Image processing

Images were processed using open-source FIJI processing software^80^. For cell counting of Sox9 and Otx2 and for images of co-staining, a single slice from the middle of the collected Z-stacks was chosen to prevent false overlap of cells in different planes. For all other cell counting and images, max intensity Z-projections were created using the full Z stack.

#### Cell Counting

Cell counting was performed on cryosectioned, immunostained, and imaged slides. All counted images were taken at 20X or 40X magnification and investigators were masked to the sample condition in all cases. Two images taken at the same magnification were counted for each sample and were independently counted by at least two investigators to ensure consistency. Images chosen for counting were taken in roughly analogous regions of the central retina, identified by the presence of the optic nerve, and taken in proximal sections on each side of the nerve. Counts per sample were taken as the average of the two images. All sample counts were normalized to the area of the retina, defined by the DAPI-positive area encompassing the outer nuclear layer, inner nuclear layer, and ganglion cell layer.

### qPCR

#### RNA Purification and cDNA Purification

Eyes from litter- and sex- matched pairs were harvested from miR-9-2 enhancer knockout animals (n=3) and retinas were dissected out. Tissue was immediately placed in 500 uL of Trizol and RNA was extracted according to standard Trizol kit protocol. Briefly, samples were incubated for 5 minutes in solution and then homogenized before chloroform extraction. To separate phases, 0.1 mL of chloroform was added to each sample, incubated 3 minutes, and centrifuged at 12000g at 4C for 15 minutes. The aqueous phase was collected and remainder of sample discarded. Following RNA extraction, 100% ethanol was added at 1:1 concentration with extraction solution. Samples were then purified using the Zymo RNA Clean and Concentrator Kit with DNAse treatment according to protocol. Samples were immediately stored at −80C until use. When ready to generate cDNA, reverse transcription was performed using the High Capacity cDNA Reverse Transcription Kit and samples were processed according to standard protocol. Samples were stored at − 80C until use.

#### qPCR

cDNA from bulk retinal tissue generated from litter- and sex-matched miR-9-2 enhancer knockout and wildtype pairs was collected (n=3). Primers were designed for *Tuba1a* (reference gene) and *Mir9-2hg* to confirm whether the putative enhancer targets miR-9-2 (Table S7). qPCR was performed using SYBR Green Master Mix according to standard protocol.

### Single Cell Preparation and Analyses

#### Tissue dissociation

Retinas were dissociated using a modified version of the 10X Genomics protocol for Nuclei Isolation from Embryonic Mouse Brain Tissue for Single Cell Multiome ATAC + Gene Expression Sequencing. Briefly, flash frozen retina tissue was submerged in lysis buffer (10mM Tris 7.4, 10mM NaCl, 3 mM MgCl2, 1% BSA, 0.1% Tween 20, 0.1% IGEPAL CA-630, 1U/uL RNAse inhibitor, and 1mM DTT if for multiome), immediately without thawing. Tissue was transferred to and broken up by glass dounce (10X dounce A, then 10X dounce B) before being transferred to clean tubes. Tissue was incubated in dissociation buffer for a total of 10 minutes (and agitated halfway through). Lysis was immediately neutralized in wash buffer (10mM Tris 7.4, 10mM NaCl, 3 mM MgCl2, 1% BSA, 0.1% Tween 20, 1U/uL RNAse inhibitor) following incubation, and nuclei were passed through a 70-micron filter and subsequent 40-micron filter (BelArt H13680-0070; BelArt H13680-0040) to clear debris. Nuclei were pelleted and washed with wash buffer twice to clear ambient RNA before being counted using Acridine Orange/Propidium Iodine Cell Stain on the ThermoFisher Automated Countess II automated cell counter. Nuclei were spun and resuspended in 1 mL of nuclei buffer (1% BSA, 1U/uL RNAse inhibitor in 1X PBS) to reach target nuclei concentration plus 10% for 10,000 cells for input into 10X Multiome or 10X 3’ GEX expression, based on 10X guidelines. To verify the final concentration, resuspended nuclei were recounted using the Countess before proceeding with 10X library generation.

#### Library preparation

For E16.5 and P0 samples, libraries were generated following the protocols for the 10X Single Cell Chromium NextGEM 3’ v3.1 kit. Equal numbers of littermate, sex-matched male and female pairs (n=2) were used to generate individual libraries. For 3-week samples, libraries were generated for sequencing following the protocols for the 10X Chromium Single Cell Multiome ATAC + Gene Expression kit. Littermate, sex-matched male and female pairs were used to generate individual libraries (n=3); due to poor sample quality, one male pair was discarded leaving three female pairs, and two male pairs for analysis.

#### Sequencing

All sequencing was performed by the Northwest Genomics Center at the University of Washington. Samples were sequenced using the NovaSeq SP100 flow cell, with parameters for all sequencing runs following 10X Genomics recommendations for cycles and sequencing depth. All data was returned in FastQ file format.

#### Single Cell Alignment

All single cell data was demultiplexed and aligned using the 10X Genomics Cellranger v9.0 Count pipeline. Following alignment, raw output matrices from Cellranger were input into the Cellbender v0.3.2 remove-background pipeline to filter data for real cells and to remove ambient RNA reads^81^. Cellbender output was assessed using provided output report, and all samples that did not pass initial Cellbender quality control (QC) using default settings were run through the pipeline again using the recommended modified learning rate of 5e-5. Filtered Cellbender output matrices that passed the provided QC metrics were used for all downstream processing.

#### Processing, clustering, and cell type identification of single cell datasets

Seurat v5 was used to perform analyses of individual and aggregated samples^82^. The filtered Cellbender output matrics were used to generate Seurat objects. Quality control was performed on individual samples and cells were filtered on several metrics, including mitochondrial percentage (<5%), number of UMIs (>500), and number of genes (>500). Individual samples of each timepoint were merged for analysis and relevant metadata (genotype, sex, batch, sample) were annotated. Each timepoint was analyzed as its own dataset. Merged objects were normalized and scaled using SCTransform^83,84^. Dimensional reduction was performed, and the Harmony package^85^ was used to integrate samples before generating Uniform Manifold Approximation and Projections (UMAPs) and clustering. Merged objects were assessed for presence of doublets using scDoubletFinder^86^. As very low numbers of doublets were found and not concentrated to any one cluster, objects were not filtered. To annotate mature retinal cell types, the 3-week dataset was annotated using the automated scMCA pipeline for mouse cell class identification^87^. For any specific cell types called by scMCA, cell identification was simplified to the major cell class containing that cell type. To annotate developmental retinal cell types, the previously published mouse atlas of retinal development^19^ was used as a reference, and cell class identities were mapped to the datasets using the Seurat FindTransferAnchors function. All automated cell type identification was validated using prior knowledge of expected cell type-specific gene expression.

#### Differential gene expression analyses

To perform differential gene expression analysis, the RNA assay of annotated Seurat objects was normalized using Seurat NormalizeData and filtered using the EdgeR^88,89^ filterbyExp (with default parameters) to remove genes with inadequate expression for statistical power. Presto^46^ was used to determine differentially expressed genes between wildtype and knockout samples for each cell type of interest. Predicted miR-9 targets in the differentially expressed genes were identified using TargetScan8.0^47,48^. Differentially expressed genes for each sample and relevant cell types were plotted as volcano plots using the EnhancedVolcano^90^ R package.

#### Pathway Analysis

To perform gene set enrichment analysis, the unfiltered list of differentially expressed genes from Presto was ranked according to the Area Under the Curve metric multiplied by the sign of logFC. We used gseGO (default settings) from ClusterProfiler^51^ to identify enriched pathways from the list of Gene Ontology Biological Processes database^52,53^ and plotted the top enriched pathways using the ClusterProfiler dotplot function.

### Quantification and Statistical Analysis

To determine enrichment of miR-9-2 targets in differentially expressed genes at timepoints and cell types of interest, predicted miR-9 targets were identified using TargetScan8.0^47,48^. To check for enrichment, hypergeometric tests were performed using base R. For bar graphs, all plots were created and analyzed with GraphPad and are shown with error bars +/- SD. Mann Whitney U tests were performed to determine whether cell counts were significantly different. For all statistical analyses, p≤0.05 were considered statistically significant.

## Data Availability

All snRNA-seq files available at GEO with accession number GSE325932. All unique code generated in this study is available at github.com/TCherryLab/miR-9-2-KO-2026.

## ACKNOWLEDGMENTS

We would like to thank all members of the Cherry Lab, CDBRM, the Lowy Medical Research Institute, as well as the LMRI Board of Scientific Governors, for helpful discussions about this project. We thank the animal care team at Seattle Children’s Research Institute for their hard work and daily maintenance of our animal colony. We would also like to thank the University of Washington Vision Core and Dr. Abbi Engel for their assistance with the TEM experiments. This work was supported by grants from the NIH National Eye Institute (R01EY028584 & R01EY033364) and the Lowy Medical Research Institute to T.J.C., an NIH P30 grant from the NIH National Eye Institute (P30EY001730) to the UW Vision Core, and an NIH T32 award (EY007031) to L.K.C. and an NIH F31 grant (EY035932) to L.K.C.

## AUTHOR CONTRIBUTIONS

L.K.C., A.J., S.X., E.D.T., and T.J.C. designed and performed experiments and analyses associated with the manuscript. L.K.C. performed bioinformatic analyses. A.J. and L.K.C. performed animal management and histology. L.K.C and T.J.C. conceptualized study design and wrote the paper with input and feedback from all co-authors.

## FIGURE LEGENDS

**Supplemental Figure 1. Quality control metrics, UMAP, and differential expression analysis for snRNA-sequencing of retinas from 5-week miR-9-2 enhancer knockout and littermate control mice (related to Fig. 1)**. **A.** Cell counts of high-quality cells from each WT and miR-9-2 enhancer KO sample for snRNA-seq. **B.** UMAPs of all enhancer knockout and WT pairs, annotated by cell class. **C.** Quality control metrics for individual snRNA-seq samples. **D.** Stacked bar plots of proportions of individual cell classes in WT and miR-9-2 enhancer knockout mice. **E.** Volcano plot of differentially expressed genes comparing KO and WT MG. Thresholds (dashed lines) correspond to p≤0.01 and FC≥10%. Wilcoxon rank sum test, Benjamini-Hochberg correction. **F.** UMAPs, split by genotype, for WT and miR-9-2 enhancer knockout retinas, annotated by cell type. Abbreviations: AC = amacrine cells, BC = bipolar cells, HC = horizontal cells, MG = Müller glia, RGC = retinal ganglion cells.

**Supplemental Figure 2. Immunofluorescent analysis of cell classes in miR-9-2 enhancer knockout and WT mice (related to Fig. 1). A.** Immunolabeling of Sox9 (green) and Otx2 (magenta) to label MG and bipolar cells, respectively, in WT and miR-9-2 enhancer knockout retinas from 5-week-old mice. **B.** Representative images of immunofluorescence for individual cell classes (green), including cones (Arr3), RGCs (Rbpms; Pax6), microglia (Iba1), and ACs (Pax6). All scale bars = 50 microns. **C.** Cell count values for individual cell types in miR-9-2 KO and WT retinas, including microglia, cells in GCL, horizontal cells, ACs, cones, RGCs, bipolar cells, and MG. Counts normalized to retinal area. Values reported as mean +/- 1SD. ANOVA, post hoc t-test, p > 0.05 for all. Abbreviations: GCL = ganglion cell layer.

**Supplemental Figure 3. UCSC Genome Browser tracks of sequencing reads from miR-9-2 knockout and wildtype animals (related to Fig. 1). A.** Representative UCSC genome browser tracks at genomic locus of miR-9-2 host gene in mouse. Blue highlight indicates miR-9 hairpin. Tracks show number of sequencing reads corresponding to each genomic position.

**Supplemental Figure 4. Quality control metrics and cell class markers for single-nucleus RNA-sequencing of miR-9-2 KO and WT animals (related to Fig. 2). A.** Table of cell counts of high-quality cells for individual WT and miR-9-2 KO samples at each profiled timepoint. **B.** Quality control metrics from snRNA-sequencing for each sample. **C.** Heatmap of annotated cell types and corresponding cell class marker expression for all E16.5 and P0 combined samples. **D.** Heatmap of annotated cell types and corresponding cell class marker expression for all 3-week samples.

**Supplemental Figure 5. snRNA-seq analysis of proportions for cell classes and cell cycle phases of miR-9-2 KO and WT retinas (related to Fig. 2). A.** Stacked bar plots representing proportions of cell types for each sample at E16.5, P0, and 3-week timepoints. **B.** Stacked bar plots of cell cycle phase proportions for WT and KO samples at each developmental timepoint. **C.** Stacked bar plots of cell cycle phases for early and late RPCs at E16.5 and P0 and developing cones at E16.5. Data fit using EdgeR generalized linear model with Benjamini-Hochberg correction^88,91^, ns for all cell classes except E16.5 Developing Cones, G2M, p≤0.05

**Supplemental Figure 6. Markers of cell death in miR-9-2 KO and WT retinas (related to Fig. 2**; **Fig.4-**6). Immunolabelling of apoptotic cells with Cas3 (magenta) in miR-9-2 KO and WT retinas at E16.5 **(A)** and **(B)** P0. **C.** Images of TUNEL assay (green) to label apoptotic cells, in miR-9-2 KO and WT retinas at 3-weeks. Abbreviations: WT = wildtype, KO = knockout, TUNEL = terminal deoxynucleotidyl transferase dUTP Nick End Labeling.

**Supplemental Figure 7 (related to Fig. 3). Expression of paralogous miR-9 host genes in individual cell classes of miR-9-2 KO and WT retinas.** snRNA-seq heatmaps of average expression for miR-9 host genes, *miR-9-1hg* (A) and *miR-9-3hg* (B), in miR-9-2 KO and WT cells of each cell class. Gray boxes indicate that the cell class was only present in negligible numbers/not present. Scale (0-20) is consistent across all heatmaps and in corresponding Figure 3F.

**Table S1. snRNA-seq differential abundance analysis of cell classes at each timepoint for miR-9-2 KO and WT retinas.**

**Table S2. snRNA-seq differential abundance analysis of cell cycle phases at each timepoint for miR-9-2 KO and WT retinas.**

**Table S3. Differential gene expression analysis from snRNA-seq for each cell type in developing E16.5 miR-9-2 KO and WT retinas.**

**Table S4. Differential gene expression analysis from snRNA-seq for each cell type in developing P0 miR-9-2 KO and WT retinas.**

**Table S5. Differential gene expression analysis from snRNA-seq for each cell type in mature 3-week miR-9-2 KO and WT retinas.**

**Table S6. Enrichment of computationally predicted microRNA targets in differentially expressed genes across cell class and timepoint in miR-9-2 KO and WT retinas.**

**Table S7. Primers generated for qPCR analysis and genotyping.**

