## Supplemental Figures for "Disease-associated microRNA, miR-9-2, regulates timing of retinal progenitor cell competence and maintenance of Müller glial identity"

**A**

| Sample | eKO1 | eKO2 | eKO3 | eWT1 | eWT2 | eWT3 |
|---|---|---|---|---|---|---|
| # Cells | 2640 | 3409 | 3141 | 3263 | 3302 | 2572 |

**B**

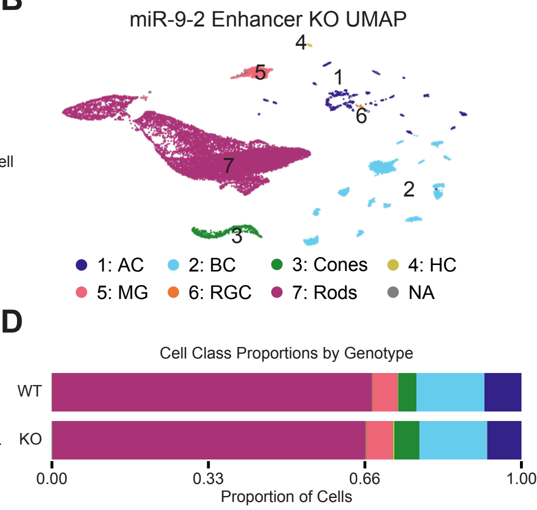


**C**

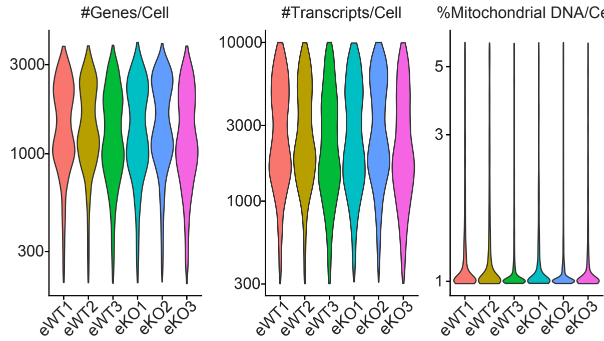


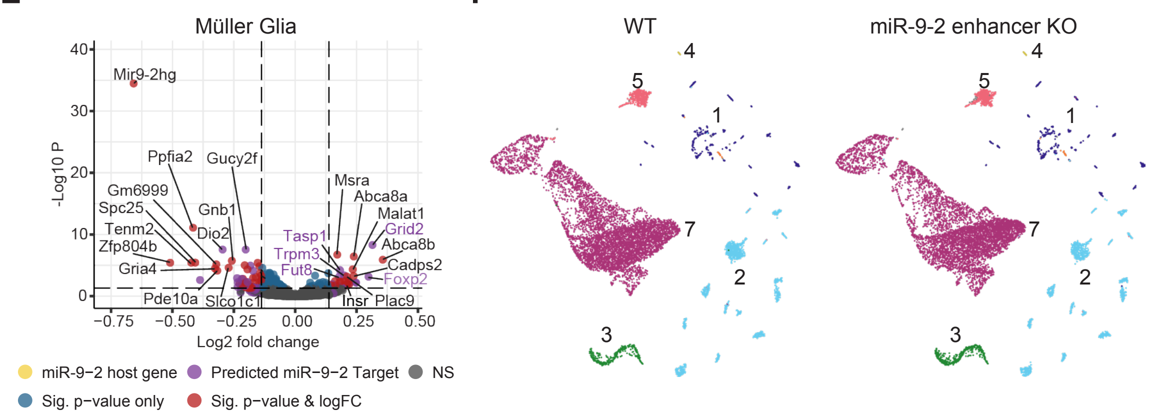


SFigure 1

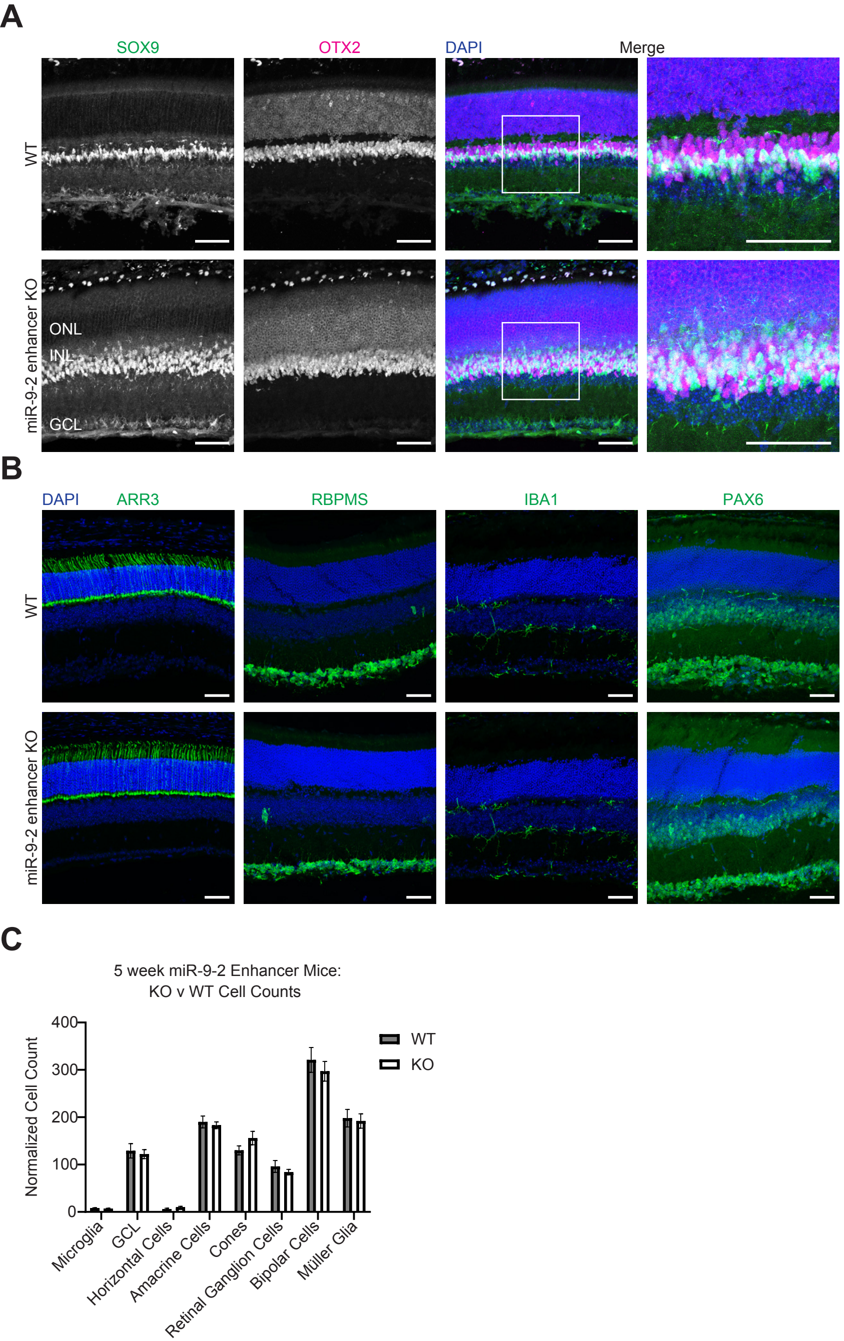
A
SOX9
OTX2
DAPI
Merge
WT
miR-9-2 enhancer KO
ONL
INL
GCL
B
DAPI
ARR3
RBPMS
IBA1
PAX6
WT
miR-9-2 enhancer KO
C
5 week miR-9-2 Enhancer Mice:
KO v WT Cell Counts
Normalized Cell Count
400
300
200
100
0
WT
KO
Microglia
GCL
Horizontal Cells
Amacrine Cells
Cones
Retinal Ganglion Cells
Bipolar Cells
Müller Glia

SFigure 2

A

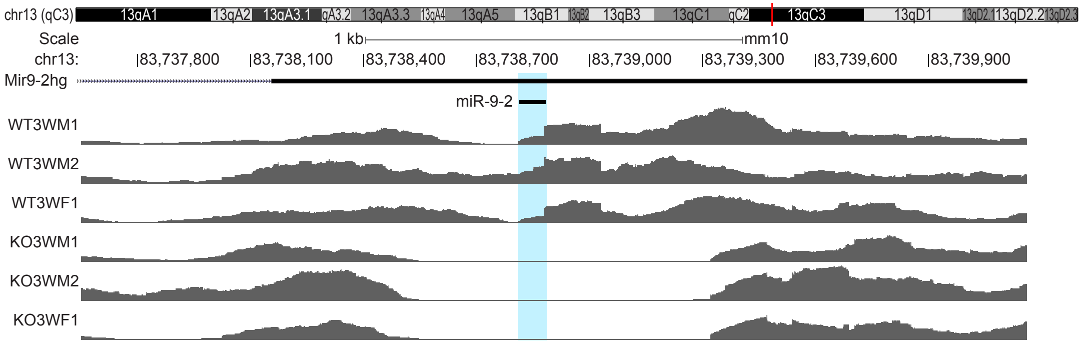
chr13 (qC3)
13qA1
13qA2
13qA3.1
qA3.2
13qA3.3
13qA4
13qA5
13qB1
13qB2
13qB3
13qC1
qC2
13qC3
13qD1
13qD2.1
13qD2.2
13qD2.3
Scale
1 kb
mm10
chr13:
83,737,800
83,738,100
83,738,400
83,738,700
83,739,000
83,739,300
83,739,600
83,739,900
Mir9-2hg
miR-9-2
WT3WM1
WT3WM2
WT3WF1
KO3WM1
KO3WM2
KO3WF1

SFigure 3

A

| Stage | Genotype | Sex | # of Cells |
|---|---|---|---|
| E16.5 | WT | F1 | 4644 |
| | | F2 | 4963 |
| | | M1 | 4236 |
| | | M2 | 4892 |
| | KO | F1 | 4284 |
| | | F2 | 5023 |
| | | M1 | 6005 |
| | | M2 | 7527 |
| P0 | WT | F1 | 5344 |
| | | F2 | 8410 |
| | | M1 | 4028 |
| | | M2 | 2798 |
| | KO | F1 | 8606 |
| | | F2 | 11251 |
| | | M1 | 2998 |
| | | M2 | 3256 |
| 3 week | WT | F1 | 6347 |
| | | F2 | 8319 |
| | | F3 | 2684 |
| | | M1 | 5785 |
| | | M3 | 4084 |
| | KO | F1 | 7998 |
| | | F2 | 3338 |
| | | F3 | 3494 |
| | | M1 | 5591 |
| | | M3 | 2073 |

B

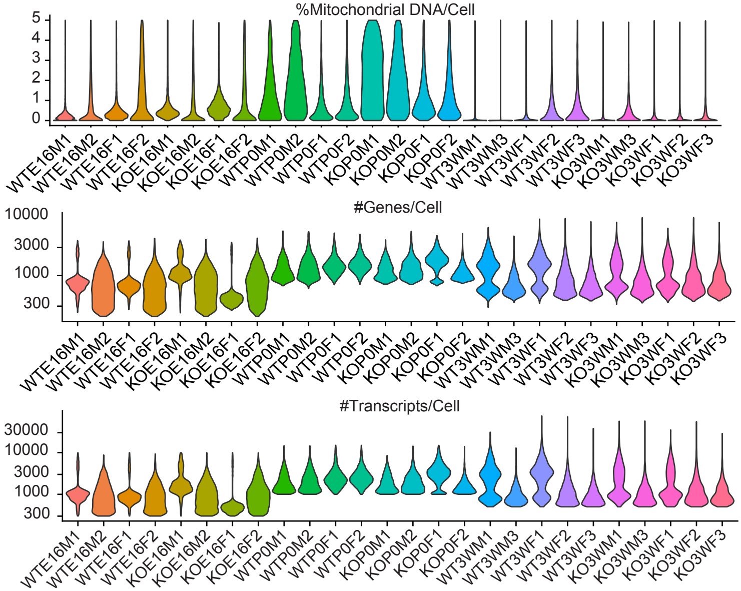


C

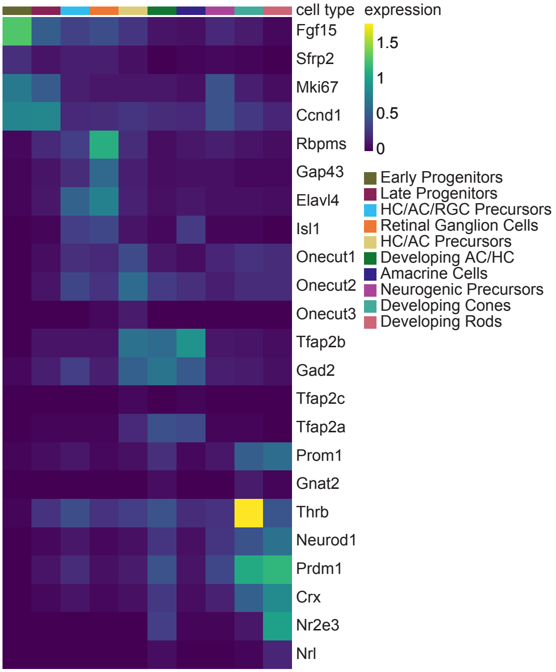


D

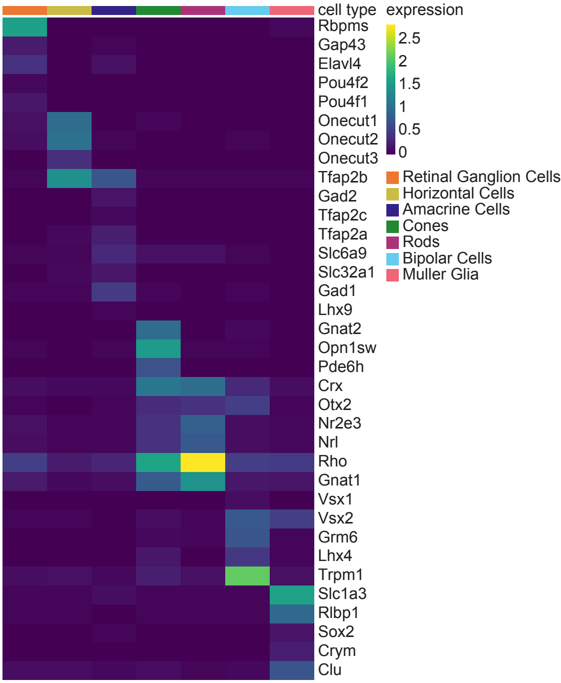


SFig 4

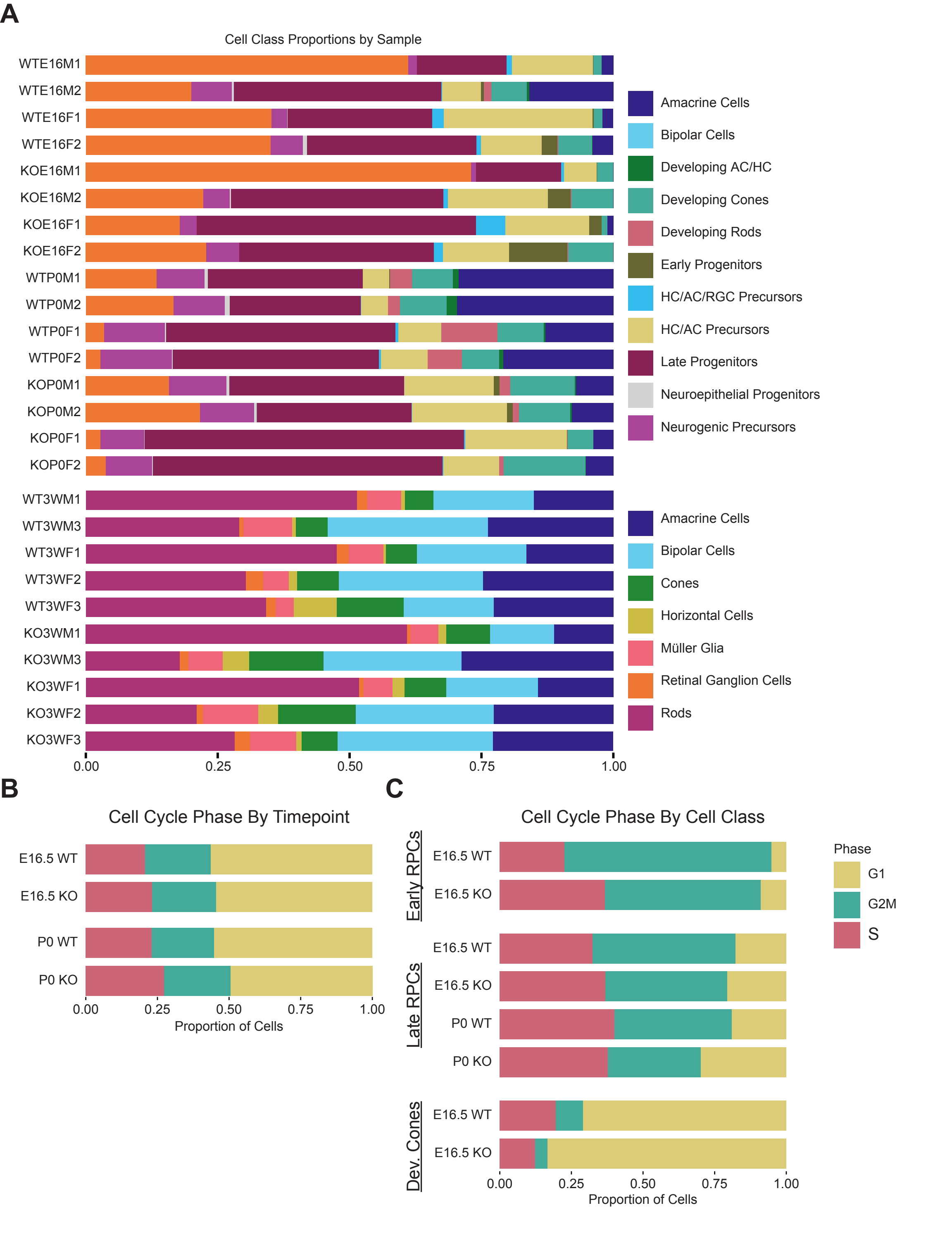
A
Cell Class Proportions by Sample
WTE16M1
WTE16M2
WTE16F1
WTE16F2
KOE16M1
KOE16M2
KOE16F1
KOE16F2
WTP0M1
WTP0M2
WTP0F1
WTP0F2
KOP0M1
KOP0M2
KOP0F1
KOP0F2
WT3WM1
WT3WM3
WT3WF1
WT3WF2
WT3WF3
KO3WM1
KO3WM3
KO3WF1
KO3WF2
KO3WF3
0.00
0.25
0.50
0.75
1.00
Amacrine Cells
Bipolar Cells
Developing AC/HC
Developing Cones
Developing Rods
Early Progenitors
HC/AC/RGC Precursors
HC/AC Precursors
Late Progenitors
Neuroepithelial Progenitors
Neurogenic Precursors
Amacrine Cells
Bipolar Cells
Cones
Horizontal Cells
Müller Glia
Retinal Ganglion Cells
Rods
B
Cell Cycle Phase By Timepoint
E16.5 WT
E16.5 KO
P0 WT
P0 KO
0.00
0.25
0.50
0.75
1.00
Proportion of Cells
C
Cell Cycle Phase By Cell Class
Early RPCs
E16.5 WT
E16.5 KO
Late RPCs
E16.5 WT
E16.5 KO
P0 WT
P0 KO
Dev. Cones
E16.5 WT
E16.5 KO
0.00
0.25
0.50
0.75
1.00
Proportion of Cells
Phase
G1
G2M
S

SFigure 5

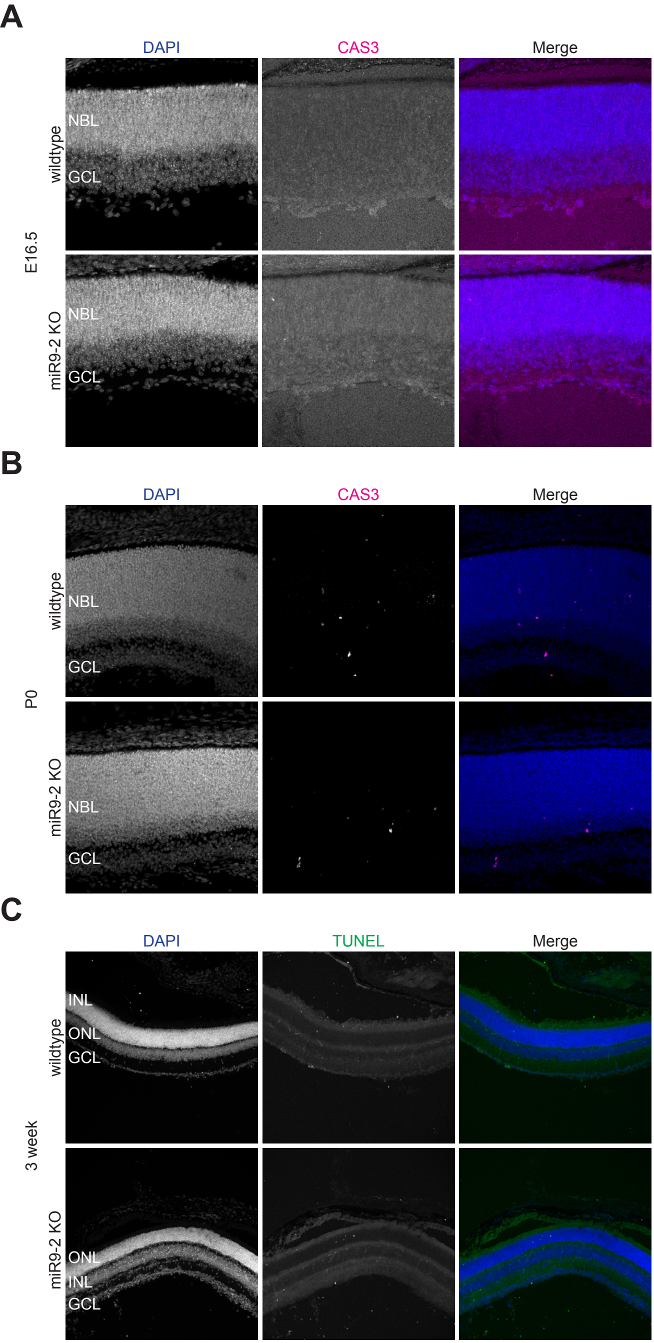
A
DAPI
CAS3
Merge
E16.5
wildtype
NBL
GCL
miR9-2 KO
NBL
GCL
B
DAPI
CAS3
Merge
P0
wildtype
NBL
GCL
miR9-2 KO
NBL
GCL
C
DAPI
TUNEL
Merge
3 week
wildtype
INL
ONL
GCL
miR9-2 KO
ONL
INL
GCL

SFigure 6

**A**

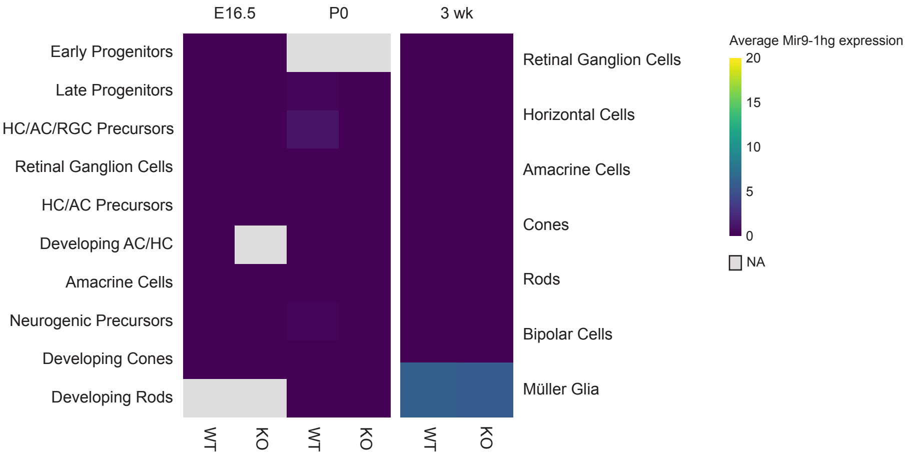
E16.5
P0
3 wk
Early Progenitors
Late Progenitors
HC/AC/RGC Precursors
Retinal Ganglion Cells
HC/AC Precursors
Developing AC/HC
Amacrine Cells
Neurogenic Precursors
Developing Cones
Developing Rods
Retinal Ganglion Cells
Horizontal Cells
Amacrine Cells
Cones
Rods
Bipolar Cells
Müller Glia
WT
KO
WT
KO
WT
KO
Average Mir9-1hg expression
20
15
10
5
0
NA

**B**

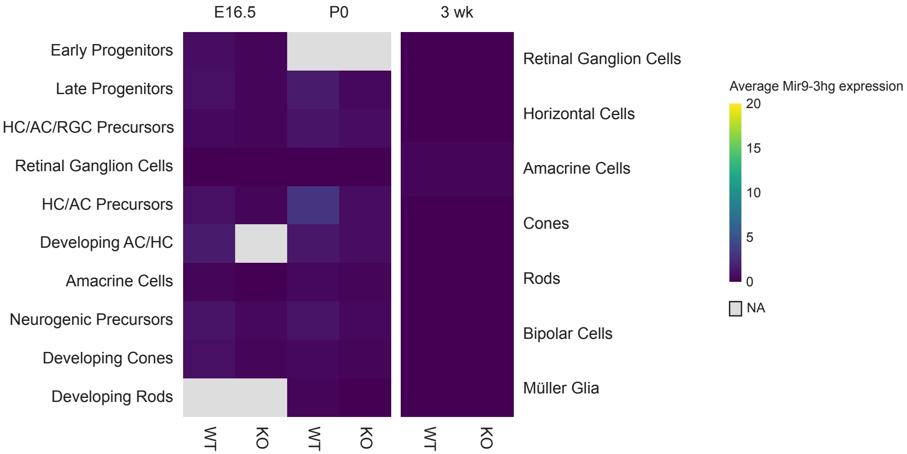
E16.5
P0
3 wk
Early Progenitors
Late Progenitors
HC/AC/RGC Precursors
Retinal Ganglion Cells
HC/AC Precursors
Developing AC/HC
Amacrine Cells
Neurogenic Precursors
Developing Cones
Developing Rods
Retinal Ganglion Cells
Horizontal Cells
Amacrine Cells
Cones
Rods
Bipolar Cells
Müller Glia
WT
KO
WT
KO
WT
KO
Average Mir9-3hg expression
20
15
10
5
0
NA

**SFigure 7**
