## Supplementary material for "Disease-associated microRNA, miR-9-2, regulates timing of retinal progenitor cell competence and maintenance of Müller glial identity": TableS1

**Supplemental Table 1**

**E16.5 Differential Abundance of Celltypes**

| **Celltype** | **logFC** | **logCPM** | **F** | **PValue** | **FDR** |
| --- | --- | --- | --- | --- | --- |
| Amacrine Cells | -4.4115 | 15.52663 | 16.97575 | 0.000806 | 0.005643 |
| Early Progenitors | 1.916568 | 15.1692 | 3.293212 | 0.089895 | 0.314631 |
| HC/AC/RGC Precursors | 0.74572 | 14.62653 | 0.828488 | 0.377328 | 0.666467 |
| HC/AC Precursors | -0.40935 | 17.88558 | 0.389238 | 0.542068 | 0.666467 |
| Neurogenic Precursors | -0.36423 | 15.94909 | 0.319263 | 0.580393 | 0.666467 |
| Developing Cones | 0.415808 | 16.19644 | 0.312284 | 0.584772 | 0.666467 |
| Late Progenitors | 0.26194 | 18.96957 | 0.192684 | 0.666467 | 0.666467 |

**P0 Differential Abundance of Celltypes**

| **Celltype** | **logFC** | **logCPM** | **F** | **PValue** | **FDR** |
| --- | --- | --- | --- | --- | --- |
| Amacrine Cells | -1.96435 | 17.1562 | 24.78963 | 0.000596 | 0.007157 |
| Developing AC/HC | -2.7723 | 12.63844 | 14.61208 | 0.00305 | 0.015729 |
| Developing_Rods | -2.62893 | 15.09864 | 13.07892 | 0.005165 | 0.015729 |
| HC AC Precursors | 1.243622 | 16.8149 | 12.78777 | 0.005243 | 0.015729 |
| Early Progenitors | 3.857765 | 11.60389 | 4.76068 | 0.048038 | 0.115292 |
| Microglia | -2.14212 | 8.957226 | 1.424527 | 0.257124 | 0.509721 |
| Late Progenitors | 0.380593 | 18.58593 | 1.148568 | 0.309633 | 0.509721 |
| Developing Cones | 0.380879 | 16.51406 | 0.993827 | 0.342883 | 0.509721 |
| HC/AC/RGC Precursors | -0.73534 | 11.18896 | 0.731719 | 0.411873 | 0.509721 |
| Neuroepithelial Progenitors | -0.71579 | 12.00721 | 0.694188 | 0.424767 | 0.509721 |

**3-Week Differential Abundance of Celltypes**

| **Celltype** | **logFC** | **logCPM** | **F** | **PValue** | **FDR** |
| --- | --- | --- | --- | --- | --- |
| Retinal Ganglion Cells | -0.83547 | 14.08346 | 3.13309 | 0.097478 | 0.781119 |
| Cones | 0.406629 | 16.4393 | 2.109844 | 0.173582 | 0.781119 |
| Bipolar Cells | -0.22487 | 17.7693 | 0.915124 | 0.358645 | 0.804438 |
| Amacrine Cells | -0.22207 | 17.61066 | 0.830883 | 0.380932 | 0.804438 |
| Horizontal Cells | 0.534959 | 14.6046 | 0.470902 | 0.504541 | 0.804438 |
| Muller Glia | 0.149446 | 16.02648 | 0.275964 | 0.609558 | 0.804438 |
| Rods | 0.131753 | 18.49265 | 0.201805 | 0.661699 | 0.804438 |
| Rgs16 | -0.26105 | 13.12049 | 0.137187 | 0.715056 | 0.804438 |
| Astrocytes | -0.15576 | 11.64168 | 0.022005 | 0.883674 | 0.883674 |
