## Supplementary material for "Disease-associated microRNA, miR-9-2, regulates timing of retinal progenitor cell competence and maintenance of Müller glial identity": TableS2

**Supplemental Table 2**

**E16.5 Differential Abundance of Cell Cycle Phases**

| **Cell Type** | **Phase** | **logFC** | **logCPM** | **F** | **P-Value** | **FDR** |
| --- | --- | --- | --- | --- | --- | --- |
| Bulk | S | 0.170609 | 17.76068 | 0.680579 | 0.430725 | 0.919256 |
| Bulk | G1 | -0.04967 | 19.08933 | 0.079391 | 0.784501 | 0.919256 |
| Bulk | G2M | -0.0459 | 17.75074 | 0.010869 | 0.919256 | 0.919256 |
| Amacrine Cells | S | 1.019454 | 17.34773 | 0.854493 | 0.376946 | 0.598356 |
| Amacrine Cells | G1 | -0.18279 | 19.58102 | 0.313857 | 0.586339 | 0.598356 |
| Amacrine Cells | G2M | 0.828249 | 15.92189 | 0.299284 | 0.598356 | 0.598356 |
| Developing Cones | G2M | -1.08275 | 16.11751 | 8.695405 | 0.008589 | 0.025767 |
| Developing Cones | S | -0.61806 | 17.1539 | 3.622992 | 0.073102 | 0.109653 |
| Developing Cones | G1 | 0.162154 | 19.60834 | 0.808012 | 0.380576 | 0.380576 |
| Early Progenitors | S | 0.43565 | 18.2369 | 0.242294 | 0.629843 | 0.847182 |
| Early Progenitors | G2M | -0.2635 | 18.88363 | 0.098579 | 0.758016 | 0.847182 |
| Early Progenitors | G1 | 0.378635 | 17.72326 | 0.038531 | 0.847182 | 0.847182 |
| HC/AC Precursors | G2M | -0.82207 | 16.2147 | 1.590259 | 0.242648 | 0.391894 |
| HC/AC Precursors | S | -0.67417 | 17.10157 | 1.458862 | 0.261263 | 0.391894 |
| HC/AC Precursors | G1 | 0.186828 | 19.58553 | 0.714572 | 0.422153 | 0.422153 |
| HC/AC/RGC Precursors | S | -0.17999 | 17.59292 | 0.189534 | 0.670363 | 0.782577 |
| HC/AC/RGC Precursors | G1 | 0.058525 | 19.42862 | 0.111591 | 0.744013 | 0.782577 |
| HC/AC/RGC Precursors | G2M | 0.087272 | 17.04157 | 0.079145 | 0.782577 | 0.782577 |
| Late Progenitors | S | 0.179067 | 18.42698 | 0.18899 | 0.671926 | 0.975105 |
| Late Progenitors | G2M | -0.12765 | 18.67359 | 0.048025 | 0.830998 | 0.975105 |
| Late Progenitors | G1 | -0.03927 | 17.81457 | 0.001031 | 0.975105 | 0.975105 |
| Neurogenic Precursors | S | 0.113722 | 17.83512 | 0.282147 | 0.609728 | 0.875764 |
| Neurogenic Precursors | G2M | -0.10641 | 18.87982 | 0.051333 | 0.826467 | 0.875764 |
| Neurogenic Precursors | G1 | 0.12088 | 18.14987 | 0.026063 | 0.875764 | 0.875764 |
| Retinal Ganglion Cells | G2M | -0.23102 | 15.59741 | 0.018648 | 0.892896 | 0.996206 |
| Retinal Ganglion Cells | G1 | 0.013261 | 19.66404 | 6.29E-05 | 0.99376 | 0.996206 |
| Retinal Ganglion Cells | S | -0.00809 | 16.88191 | 2.32E-05 | 0.996206 | 0.996206 |

**P0 Differential Abundance of Cell Cycle Phases**

| **Cell Type** | **Phase** | **logFC** | **logCPM** | **F** | **P-Value** | **FDR** |
| --- | --- | --- | --- | --- | --- | --- |
| Bulk | S | 0.186462 | 17.76343 | 0.309896 | 0.584599 | 0.685318 |
| Bulk | G1 | -0.14491 | 19.00322 | 0.190513 | 0.66768 | 0.685318 |
| Bulk | G2M | 0.138133 | 17.94558 | 0.169614 | 0.685318 | 0.685318 |
| Amacrine Cells | S | 0.26337 | 16.8915 | 0.136394 | 0.716201 | 0.94779 |
| Amacrine Cells | G2M | 0.186861 | 15.78291 | 0.061856 | 0.806401 | 0.94779 |
| Amacrine Cells | G1 | -0.04552 | 19.65501 | 0.004409 | 0.94779 | 0.94779 |
| Developing Cones | G2M | -0.53711 | 17.60704 | 0.398444 | 0.536447 | 0.875939 |
| Developing Cones | S | 0.261803 | 17.41459 | 0.164648 | 0.690068 | 0.875939 |
| Developing Cones | G1 | 0.096617 | 19.26054 | 0.025132 | 0.875939 | 0.875939 |
| Developing Rods | S | 0.511037 | 16.79744 | 0.398403 | 0.535849 | 0.843685 |
| Developing Rods | G2M | 0.6221 | 15.39539 | 0.348222 | 0.562457 | 0.843685 |
| Developing Rods | G1 | -0.05099 | 19.68924 | 0.006115 | 0.938534 | 0.938534 |
| Developing AC/HC | G2M | -1.90425 | 17.18725 | 1.879304 | 0.187266 | 0.561797 |
| Developing AC/HC | G1 | 0.17062 | 19.49791 | 0.131101 | 0.721509 | 0.979318 |
| Developing AC/HC | S | 0.022937 | 17.53316 | 0.000691 | 0.979318 | 0.979318 |
| HC/AC Precursors | S | 0.062501 | 17.08873 | 0.015267 | 0.904716 | 0.983592 |
| HC/AC Precursors | G2M | 0.024621 | 16.12835 | 0.000969 | 0.97591 | 0.983592 |
| HC/AC Precursors | G1 | -0.00466 | 19.59652 | 0.00045 | 0.983592 | 0.983592 |
| Late Progenitors | G1 | 0.556014 | 17.29696 | 0.20431 | 0.663081 | 0.828333 |
| Late Progenitors | G2M | -0.1326 | 18.91059 | 0.071551 | 0.795769 | 0.828333 |
| Late Progenitors | S | -0.06938 | 18.40594 | 0.050155 | 0.828333 | 0.828333 |
| Neurogenic Precursors | G2M | 0.187585 | 18.58777 | 0.119042 | 0.738964 | 0.885445 |
| Neurogenic Precursors | G1 | -0.2161 | 18.48396 | 0.112824 | 0.745585 | 0.885445 |
| Neurogenic Precursors | S | 0.033374 | 17.88913 | 0.022123 | 0.885445 | 0.885445 |
| Retinal Ganglion Cells | S | 1.20975 | 15.72274 | 1.31289 | 0.266878 | 0.541064 |
| Retinal Ganglion Cells | G2M | 0.996581 | 15.46408 | 0.879663 | 0.360709 | 0.541064 |
| Retinal Ganglion Cells | G1 | -0.10714 | 19.786 | 0.013563 | 0.908578 | 0.908578 |
