## Supplementary material for "Disease-associated microRNA, miR-9-2, regulates timing of retinal progenitor cell competence and maintenance of Müller glial identity": TableS7

**Supplemental Table 7**

Primers for qPCR, Related to STAR Methods, Figure 1

| **Gene** | **Forward Primer** | **Reverse Primer** |
| --- | --- | --- |
| Tuba1a control | ggcagtgttcgtagacctggaa | ctccttgccaatggtgtagtgg |
| miR9-2-hg | TGGAGTTCAGCCAGAGGAAG | TCGGTGACCTTGAAGGAGTT |

Primers for genotyping, Related to STAR Methods

| **Gene** | **Forward Primer** | **Reverse Primer** |
| --- | --- | --- |
| miR9-2-lacZ-fl-neo | GAGGGGCTTGTACAACTGGA | CCCTTTTCCCTGCTAGCTCT |
| miR9-2 KO | ACATCTCAGTAGAGGAAGGC | CGTGGGTTGTGGCGTAAA |
|  | GGGTGACCACTGATGCATAT |  |
| miR9-2 Enh KO | CAATTGAAGGAGGAGTGTGCTTTA | GACAGTTGTCATGGACATCACTG |
|  | GAGCAGAGTTAATCTGGGAC |  |
| Jarid1c/d | CTGAAGCTTTTGGCTTTGAG | CCACTGCCAAATTCTTTGG |
| FlpE | GCTGAACTAACCTATTTATGTTGGATG | GCGTCTTGTATTTAAACTGGAGTG |
|  | CACGTGGGCTCCAGCATT | TCACCAGTCATTTCTGCCTTTG |
